# Statistical learning and uncommon soil microbiota explain biogeochemical responses after wildfire

**DOI:** 10.1101/2022.02.06.479310

**Authors:** Alexander S. Honeyman, Timothy S. Fegel, Henry F. Peel, Nicole A. Masters, David C. Vuono, William Kleiber, Charles C. Rhoades, John R. Spear

## Abstract

Wildfires are a perennial event globally and the biogeochemical underpinnings of soil responses at relevant spatial and temporal scales are unclear. Soil biogeochemical processes regulate plant growth and nutrient losses that affect water quality, yet the response of soil after variable intensity fire is difficult to explain and predict. To address this issue, we examined two wildfires in Colorado, USA across the first and second post-fire years and leveraged Statistical Learning (SL) to predict and explain biogeochemical responses. We found that SL predicts biogeochemical responses in soil after wildfire with surprising accuracy. Of the 13 biogeochemical analytes analyzed in this study, 9 are best explained with a hybrid microbiome + biogeochemical SL model. Biogeochemical-only models best explain 3 features, and 1 feature is explained equally well with hybrid or biogeochemical-only models. In some cases, microbiome-only SL models are also effective (such as predicting NH_4_^+^). Whenever a microbiome component is employed, selected features always involve uncommon soil microbiota (i.e., the ‘rare biosphere’, existing at *<* 1% relative abundance). Here, we demonstrate that SL paired with DNA sequence and biogeochemical data predict environmental features in post-fire soils, though this approach could likely be applied to any biogeochemical system.

## 2 Introduction

Wildfire is a natural event, but wildfire extent, frequency and severity are changing as a consequence of climate, fuel profiles and sources of ignition [1– 3]. While wildfire effects on ecosystems are typically assessed based on their macroscopic effects (e.g., burned trees and basin water quality impacts), the importance of the soil microbial world to ecosystem biogeochemical function is receiving increasing attention [4, 5]. The severity of wildfire substantially affects soil ecosystems inclusive of the microbiome and associated organic and inorganic chemical environments [6, 7]. Many factors including spatial heterogeneity (in soil types, vegetation, topography, etc.), soil burn severity (SBS; as High, Moderate, Low, and No Burn), and time since fire create soil complexity [8–12], so a general response to wildfire is implausible [13, 14]. However, improved predictions of post-fire microbial and biogeochemical responses will advance understanding of ecosystem disturbance and recovery. These predictions should be quantitative rather than descriptive —and will rely upon the truism that the microbial and biogeochemical worlds are intertwined [15–20]. There are post-fire legacy effects in soil [9, 21–24]—and it would be valuable to predict these long term effects with quantitative precision. However, to date, quantitative predictions about biogeochemical responses in disturbed soil remain elusive.

Generating vast amounts of data from environmental DNA and RNA has become relatively straight-forward [25–27]. Nonetheless, explaining a complicated system such as soil remains challenging—especially so in relating the microbiome to biogeochemistry at field scale. One response to these challenges is to systemically probe mechanisms with experimental designs that either rule-in or rule-out specific behavior as important. Such experiments, however, potentially introduce confirmation bias; when we look for specific behavior, we tend to find it (even if it has unknown relevance at the spatial scales that fires burn on). Further, it is unusual to uncover new, unexpected, relationships in data when we are not explicitly searching for or testing them in the first place. Rather, as suggested by Nurse in 2021—we may need to first consider ‘the flow of information through living systems,’ which in turn could help us ‘make better sense of the flood of biological data’ [28].

Soils are rich in data: 1 g contains on the order of 1 billion microbial cells [29], each of which has a genome with millions of nucleic acid base pairs. Conservatively, there is on the order of 1 petabyte (PB) of genetic information in a single gram of soil. SL and Machine Learning (ML) are increasingly being applied to gain inference from and to predict complicated behavior and patterns from large datasets [30]. Advancements in statistical theory [31] and the development of more efficient algorithms [32] have helped. The biomedical sciences [33–35], among others, are increasingly capitalizing on SL and ML techniques for a variety of health applications. Yet, despite the enormous potential of SL and ML [36], they have not been widely adopted in designed field-based environmental research [37, 38]. As such, the environmental sciences remain ripe for a wider adoption of data science and statistics [38] as our ability to generate large environmental datasets—hundreds to thousands of observations entailing thousands to millions of features (variables), i.e., ‘Big Data’—is outstripping our capacity to make sense of the inherently complicated environment at scales of importance.

To be used effectively, the dimensions of datasets need to be created at the outset with SL and ML techniques in mind as the evaluative framework. Some ML algorithms are data-hungry —requiring far more samples (*N*) than can be reliably collected at field scale. SL, on the other hand, does not require a large *N*, but modern SL machinery are required when the number of features (*p*) far exceeds *N* (the system is overdetermined). Environmental biogeochemistry is primed for SL approaches given that *p* is generally far larger than *N*.

In this application of SL to fire disturbed soils, we track soil biogeochemical recovery at two fires in Colorado, USA—the 416 Fire outside of Durango, CO and the Decker Fire outside of Salida, CO—with 47 unique spatial plots (resampled over three years and three seasons) representing a gradient of SBS classes, and both organic / mineral horizons (Org/Min-horizons) over multiple depths (Supplementary Figs. 1-3). To do this, we acquired a large microbial-biogeochemical dataset by sequencing 542 microbiome samples and quantifying the biogeochemistry of 178 samples from the same soils. There are 18 biogeochemical features that were measured in every soil sample (Supplementary Tables 1 - 4). Thirteen of these features represent non-redundant information and were used to train models (Supplementary Table 5). We then paired these 13 independent biogeochemical features with tens of thousands of microbiome features —quantified by the relative abundance of SSU rRNA genes as amplicon sequence variants (ASVs) via high-throughput DNA sequencing. Thus, there are on the order of 15 million data points in our study. These data represent great variance (Fig. 1) and encompass key SBS, spatial heterogeneity, time since fire, and soil horizon variables. Our aim was to address two questions with these data: 1) Is it possible to train statistical models that well-explain biogeochemical responses after wildfire?; and 2) what microbial and biogeochemical features are identified as skillful to those predictions?

**Fig. 1.**
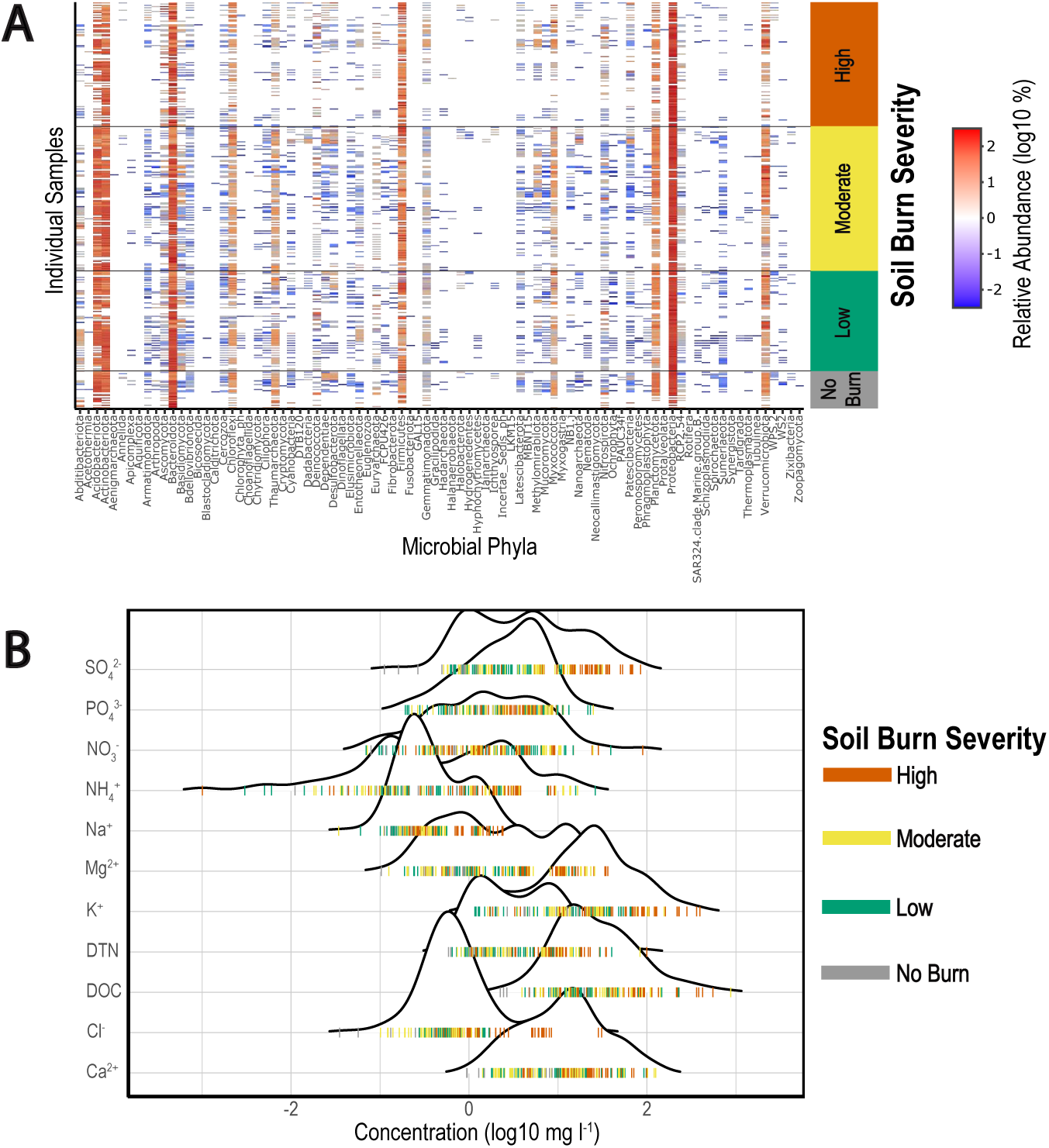
The total sample set (N_microbiome_ = 542, N_biogeochemistry_ = 178) is highly variable across soil burn severity (SBS), space, time since fire, and soil horizon. SBS, specifically, is insufficient to clearly differentiate soil samples. Due to the large number of features measured from the microbiome there are on the order of 15 million datapoints in our study. (A) The relative abundance of all detected Phyla — the highest, and simplest, level of microbial taxonomy — arranged vertically by SBS across all soil samples. The color of individual bars denotes percent relative abundance (legend at far right). Contaminant, Chloroplast and Mitochondria sequences were removed from the dataset. Although Arthropoda (and other non-microbial Eukarya) are not part of the microbiome, these DNA sequences represent real information from soil and were retained in the dataset. These non-microbial reads are few in number; for example, Arthropoda sequences have a mean relative abundance of only 0.08%. A Phyla with no detected abundance is plotted as white. A similar graphic depicting all amplicon sequence variants (ASVs) that were measured and incorporated into the study would be hundreds of times wider. In general, there are several Phyla that are common in soil—and, heuristically, these are identified by dark red bands that traverse vertically up the entire figure. Though highly abundant, these Phyla are less helpful for making predictions about soil due to their ubiquity. Rare events, i.e., bands of any color surrounded by white—are far more unique and diagnostic in predicting biogeochemical responses. (B) Measured water-extractable biogeochemical concentrations vary over several orders of magnitude in the study. Each band below the distributions is one measured extract from soil and is color-coded by SBS.

We find that sparsity-inducing linear statistical models (i.e., models that select which features are important) are capable of predicting concentrations of biogeochemical analytes in soil. We discovered which microbiological and biogeochemical features are important by *shrinking* the vast majority of coefficients to *exactly* zero using the least absolute shrinkage and selection operator (lasso) penalty applied to multiple linear regression (Equation 1) [31].

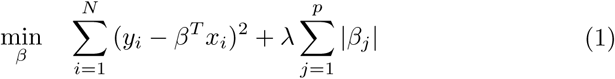

Equation (1) minimizes the squared error of predictions while the coefficients *β* are subject to a ‘penalty’ for their inclusion in the model. This minimization occurs over the target value (*y*_*i*_) and feature space (*x*_*i*_) for all samples (*i* = 1 to *N*) using all features for each sample (*j* = 1 to *p*). We estimate the weight *λ* by minimizing Equation (1) for a sequence of possible values of *λ*, and choose the one for which cross-validation predictions on held-out testing data are the most accurate. For the optimal *λ*, we identify the coefficients *β* that minimize Equation (1). The result is a set of coefficients *β* that construct the best possible linear model for predictions.

Although the conventional wisdom is that the abundant microbiota are the main drivers of ecological response [39], our research has led us to conclude that the microbial features useful for the *prediction* of responses tend to be members of the ‘rare biosphere’ and are uncommon (taxa binned at any phylogenetic rank present at *<* 1% mean relative abundance across all samples in the dataset). This suggests that abundant microbiota, though likely still major drivers in ecosystems, may be less predictive than previously understood for biogeochemical responses at field scale.

The analysis presented in this paper is intended to showcase the usefulness of SL in environmental biogeochemistry while simultaneously inviting further analysis of our dataset by the scientific community (see Data Availability section).

## 3 Methods

### Wildfire Sampling

Two wildfires in Colorado, USA were studied with recurrent sampling of 47 unique plots over the first and second post-fire years (collections from one fall, one spring, and three summers); in total, these data are 178 bulk soil collections and 542 microbiome samples. The 416 Fire, *∼* 13 miles north of Durango, Colorado in the San Juan National Forest (ignition on June 1^st^, 2018; 54,130 acres final footprint) and the Decker Fire, south of Salida, Colorado in the Rio Grande and San Isabel National Forests (detected on September 8^th^, 2019; 8,910 acres final footprint) both burned in mosaic patterns with varying SBS [40, 41]. Both fires burned in montane western forests at *∼* 8,000 ft elevation with mixed tree communities comprised of conifers (e.g., Ponderosa pine and Douglas fir) and hardwoods (e.g., quaking aspen and Gambel oak). The 416 Fire was sampled three times: June 27^th^, 2018; June 13^th^, 2019; and June 18^th^, 2020. The Decker Fire was also sampled three times: November 7^th^, 2019; May 22^nd^, 2020; and July 29^th^, 2020.

Regions of the 416 and Decker fires that represented a transect of SBS classes were targeted for sampling. 47 unique spatial plots were established (17 at the 416 Fire and 30 at the Decker Fire). Both Org-horizon and Min-horizon bulk soil samples were collected at the 416 Fire and only Org-horizon samples were collected at the Decker Fire. Biological triplicate soil samples were taken for DNA from each bulk soil collection, and the remainder bulk soil (singlicate) was saved for biogeochemistry. A detailed description of field sampling methods is provided in the Supplementary Information.

### Biogeochemistry

Bulk soil samples were air-dried, protected from light, in the humidity-controlled laboratory at 22° C for up to 14 days until mass loss from evaporation ceased. Triplicate leachate tubes were made from each sample with a 1:10 mass ratio of soil to milliQ purified water (*>*18.2 MΩ-cm resistivity) and gently mixed for 2 hours on a tabletop shaker. A bell-jar vacuum filtration system (KIMBLE®, DWK Life Sciences, NJ, USA) was used to filter leachates through two different filter types: 1) 0.7 μm glass fiber filters without binders (MilliporeSigma™ AP4004705) for dissolved organic carbon and dissolved total nitrogen (DOC/DTN) analyses; and 2) 0.45 μm PVDF membrane filters (MilliporeSigma™ Durapore™ HVLP04700) for ion chromatography (IC) analyses. The third leachate was saved, unfiltered, at 4° C for acid neutralizing capacity (ANC) and pH determination. DOC/DTN and IC filtrates were stored at -20° C. For every 20 samples filtered, raw milliQ water and a filtrate blank for each filter type were processed the same as all other samples. Laboratory analytical methods for the characterization of filtrates are reported in the Supplementary Information.

### DNA Extraction and Sequencing

#### Extraction

DNA from samples stored in Zymo ZR BashingBead™ Lysis Tubes was extracted according to manufacturer instructions for the ZymoBIOMICS™ DNA Miniprep Kit, including the optional polymerase chain reaction (PCR) inhibitor removal step (polyphenolics, humic/fulvic acids, and melanin), and then stored at -80° C. *Sequencing*. Singlicate 50 μL PCR reactions amplified the V4-5 hypervariable regions of the SSU rRNA gene with universal primers (Bacteria, Archaea, and Eukarya) [42]. Primers and reaction concentrations are reported in the Supplementary Information. Sequencing was performed with MiSeq V2 Paired End 2×250 Chemistry (Illumina Inc., San Diego, CA) at the Duke Center for Genomic and Computational Biology or the University of Colorado Anschutz Medical Campus Genomics and Microarray Core.

### Digital PCR

90 DNA samples were examined for absolute Bacterial, Archaeal, and microbial Eukaryotic abundance with a QIAcuity One 5-plex digital PCR (dPCR) instrument (Qiagen Inc. USA, Germantown, MD). dPCR samples represented both the 416 Fire —across SBS, space, time, and soil horizon —and the Decker Fire —across SBS, space, and time. A QIAcuity 8.5k 96-well Nanoplate and the QIAcuity EG PCR Kit was used for amplification and imaging according to manufacturer instructions. Primer, reaction, thermocycling, and imaging / quantification details are reported in the Supplementary Information.

### Bioinformatics and Statistical Learning

#### Bioinformatics

ASVs were called and taxonomy was assigned by Dada2 [26, 43, 44]. ASV, taxonomy, and associated sample metadata were organized as phyloseq objects [45]. Contaminating DNA reads were identified and removed by Decontam [46]. Relative abundances of taxa were computed by ampvis2 [47]. Details for each bioinformatic step are presented in the Supplementary Information.

#### Statistical Learning

All SL and statistics were conducted in R [48]. Training (75% of data) and testing datasets (the remaining 25%) were randomized from the total dataset (N_microbiome_ = 542, N_biogeochemistry_ = 178), and the process of splitting data into various feature and target matrices was automated with in-house code; within our code base, any combination of biogeochemical or microbiome data —at any user-defined phylogenetic rank —can be coerced into training, testing, feature, and target matrices immediately compatible with either of the two following statistical models. *Lasso-penalized linear regression*: The penalty weight *λ* was chosen with 10-fold cross validation on training data with the glmnet R package [32]. Root mean squared error (RMSE) was reported for both training and testing data. *Lasso-penalized principal component regression*: Principal components were computed on the full sample set in R with prcomp (options to center and scale data were set to TRUE). Lasso-penalized multiple linear regression was trained on principal components with samples from the training dataset, and cross-validation was conducted exactly as above. The testing dataset was used for plotting and model evaluation. *Randomized Training and Testing Data Splits* : For DOC and DTN predictions, model sensitivity to training / testing data randomization was assessed by training on 10 different random splits of data; each one of the random splits still received 10-fold cross-validation on training data. Median results from the 10 different data splits are reported for DOC and DTN in Table 1; plotted points (Fig. 2 & Fig. 3) are from the train/test split that produced a hybrid model testing RMSE closest to the median score. The ammonium (NH_4_^+^) case-study (Fig. 7) and all other model scores reported in the Supplementary Information use this same train/test data split. Testing the variation in NH_4_^+^ concentrations across SBS was conducted via one-way ANOVA in R with the aov() function [48].

**Table 1.**
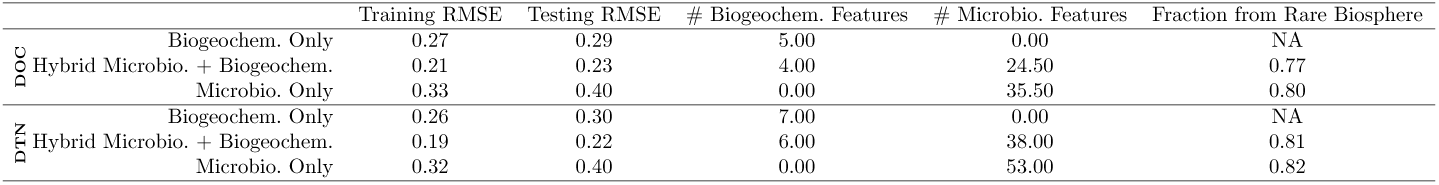
Predicting leachable dissolved organic carbon (DOC) and dissolved total nitrogen (DTN) from burned soil. DOC and DTN concentrations were log10 transformed prior to modeling. Values are median model scores across ten different training and testing dataset splits of 75% and 25% of total data, respectively. Reported numbers of features are the numbers of features selected by SL as important. A feature count of one-half is a result of calculating medians. 10-fold cross validation was also employed on each of the training sets. Different types of microbiome / biogeochemical datasets are compared. For both DOC and DTN, the hybrid biogeochemical + genera microbiome dataset performs best. Models are evaluated with the RMSE of residuals (log10-space mg l^-1^). For both model types using microbiome data, the vast majority of important genera were rare (*<* 1% mean relative abundance across all samples in the dataset).

**Fig. 2.**
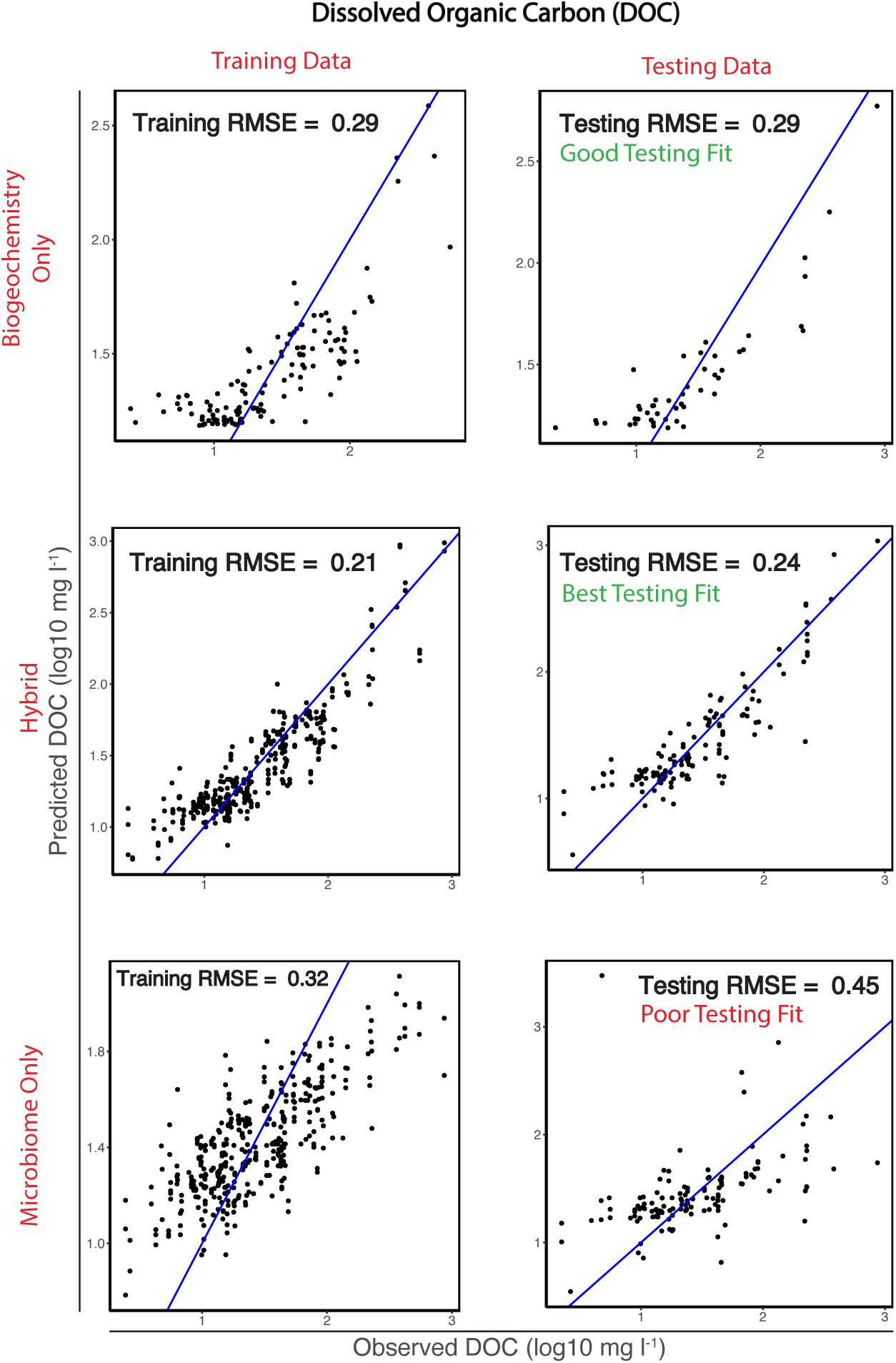
SL predictions of leachable dissolved organic carbon (DOC) from soil after wildfire. Target values were log10 transformed prior to modeling. Blue lines are a reference 1:1 line for the predicted vs. observed values (log10 mg l^-1^) —and the visual spread of data about the blue line is an important measure of model accuracy; median RMSEs (Table 1) are a fair assessment of the aggregate accuracy of a model type, but single model RMSEs (as are necessary here for plotting) are sensitive to outlier predictions. Three model types are compared: biogeochemistry alone, a hybrid biogeochemistry + genera microbiome dataset, and the genera microbiome alone. Independent training and testing datasets evaluate the accuracy of learned models. Note that axes scales are different for each scatter plot. The hybrid model performs better than either biogeochemistry or the microbiome models, alone.

**Fig. 3.**
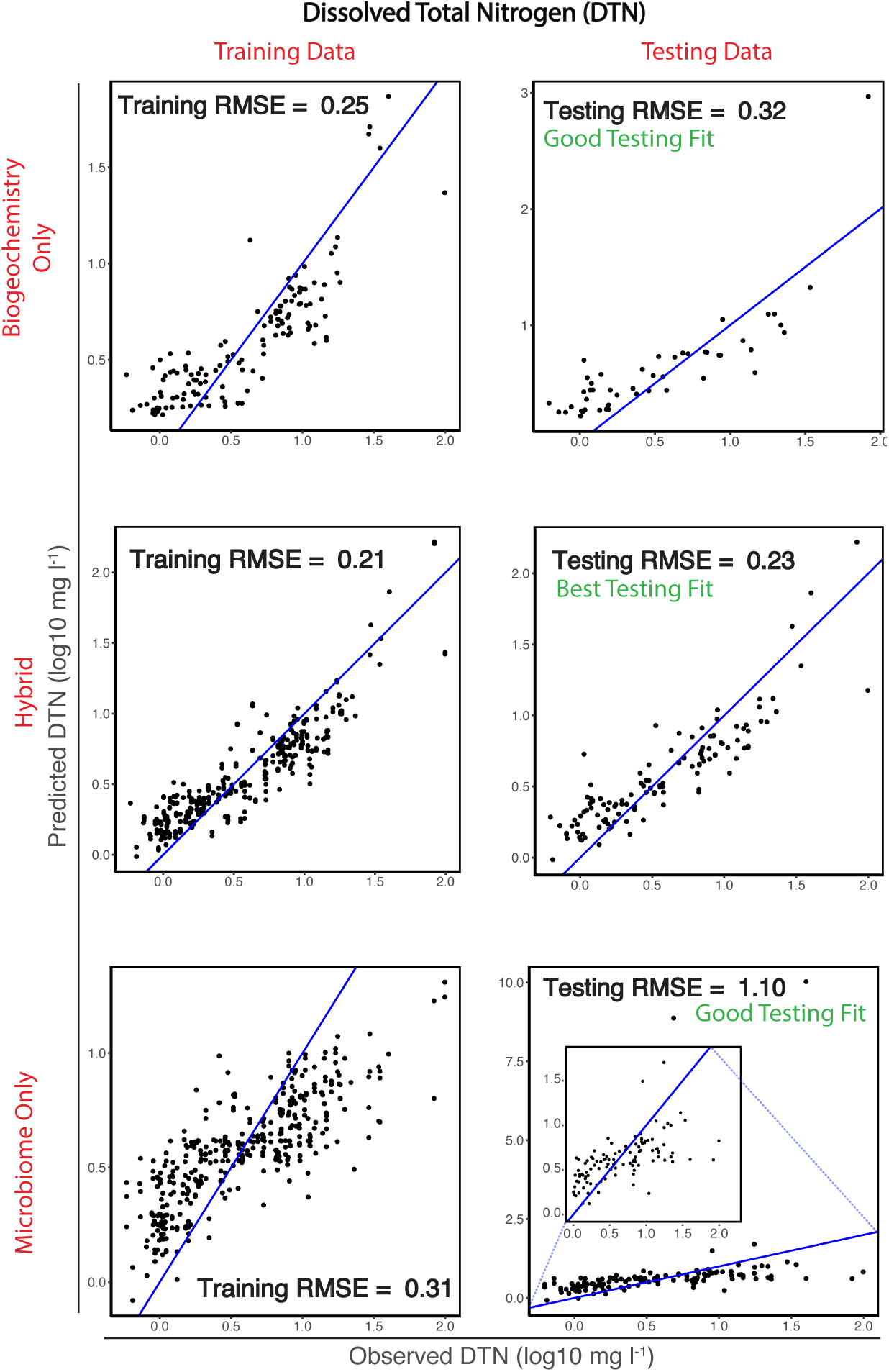
SL predictions of leachable dissolved total nitrogen (DTN) from soil after wildfire. Target values were log10 transformed prior to modeling. Blue lines are a reference 1:1 line for the predicted vs. observed values (log10 mg l^-1^) —and the visual spread of data about the blue line is an important measure of model accuracy; median RMSEs (Table 1) are a fair assessment of the aggregate accuracy of a model type, but single model RMSEs (as are necessary here for plotting) are sensitive to outlier predictions. Three model types are compared: biogeochemistry alone, a hybrid biogeochemistry + genera microbiome dataset, and the genera microbiome alone. Independent training and testing datasets evaluate the accuracy of learned models. Note that axes scales are different for each scatter plot. There are two outlier points that lengthen the y-axis in the microbiome-only testing data scatterplot; the inset scatter plot is a magnification of data around the 1:1 reference line. The hybrid model performs better than either biogeochemistry or the microbiome models, alone.

## 4 Results

### Summary of SoilBiogeochemistry

Soil undergoes obvious, rapid, transformation after fire (Supplementary Fig. 3), and there is great variance in biogeochemical profiles post-fire (Fig. 1). We observe that soil biogeochemistry is dependent on SBS, soil horizon, and time since fire and the most change in soil occurs in the first year after burning (Supplementary Tables 1-5). For example, in Org-horizons from high SBS plots at the 416 Fire, mean water-extractable dissolved organic carbon (DOC), dissolved total nitrogen (DTN), and ammonium (NH_4_^+^) decrease by 93%, 77%, and 44% from immediately post-fire to one year later (2018 to 2019), respectively (Supplementary Tables 1 - 2). Heat from fire has a limited distance that it can transfer in soil —and this distance will differ depending on the severity with which soil burned and the nature of the soil (e.g., bulk density and water content); thus, different patches of soil and different soil horizons within those patches will respond differently to wildfire. Embracing the variability of soil across principal compositional features (e.g., SBS, space, time, and soil horizon) in our study was critical to establishing a representative dataset.

### Summary of the Soil Microbiome

Several phyla of microbiota are common in burned and unburned soils (Fig. 1)—though these groups of microorganisms are not unexpected for any soil habitat: Acidobacteriota, Actinobacteriota, Bacteroidota, Thaumarchaeota, Firmicutes, Proteobacteria, and Verrucomicrobiota. There are also dozens of rare phyla present in subsets of our samples and, similarly, tens of thousands of rare amplicon sequence variants (ASVs). Despite the difficulty with which DNA can be extracted from soils with a dramatic reduction in microbial abundance—that likely also have high concentrations of PCR inhibitors—we were able to successfully amplify DNA from each SBS and reliably remove contaminating DNA sequence reads (Supplementary Fig. 5). SBS, space, time since fire, and soil horizon all impact microbial diversity as well as the amount of abundant vs. rare microbiota —as quantified by observed unique ASVs and Hill numbers [49, 50] (Supplementary Fig. 6). Soil is not sterilized after wildfire, or at least repopulates quickly. In the first post-fire sampling (weeks after burning), high SBS Org-horizon soils from both the 416 and Decker Fires contain *>* 1×10^5^, *>* 1×10^4^, and *>* 1×10^4^ copies of bacterial, archaeal, and eukaryal marker genes, respectively, per gram of soil as determined by absolute quantification via digital polymerase chain reaction (dPCR) (Supplementary Fig. 4). Within one year of burning, microbial populations (from all three domains of life) in these high SBS Org-horizons increase by at least one order of magnitude. These observed microbial recoveries are likely attributed to colonization from residual Org-horizon, upward movement from mineral and rhizosphere soils, and precipitation [51] / dust inputs.

### Statistical Learning

Our goal is to explain the complex variation represented in Fig. 1 in a way that is useful. Trained linear models with a lasso penalty term are able to successfully predict biogeochemistry concentrations —as evaluated by the root-mean-squared-error (RMSE) of residuals (a smaller RMSE means a model is more accurate) (Table 1, Fig. 2 & Fig. 3). Hybrid biogeochemistry + microbiome models are the most accurate for 9 of 13 measured biogeochemical analytes in fire-impacted soil: DTN, DOC, acid neutralizing capacity (ANC), PO_4_^3-^, SO_4_^2-^, NH_4_^+^, Na^+^, Cl^-^, and pH (Supplementary Tables 7 & 8). For NO_3_^-^, Ca^2+^, and Mg^2+^, the biogeochemical-only models are most accurate (Supplementary Table 6). K^+^ is predicted equally well with biogeochemical-only and hybrid models (Supplementary Tables 6-8). Microbiome-only models never had the best RMSE for an analyte —though predictions are still accurate beyond expectation (Supplementary Tables 9 & 10). In a direct comparison of biogeochemical-only models and microbiome-only models, the biogeochemical models are more accurate for 10 of the 13 analytes; however, for the remaining 3 analytes, PO_4_^3-^ is explained equally well by both model types, and the microbiome model is better than the biogeochemical model at predicting NH_4_^+^ and pH. We ran *>* 150 models that predict biogeochemical responses (Supplementary Tables 6 - 10); here, we provide two in-depth examples of the application of SL to microbiome + biogeochemistry datasets: 1) DOC and DTN requiring hybrid (microbiome + biogeochemistry) models; and 2) NH_4_^+^, an in-depth extension of the microbiome-only model motivated by our finding that the microbiome is better at predicting NH_4_^+^ than biogeochemistry. These two examples provide a snap-shot of the predictability of the variation in Fig. 1, but analogous in-depth explanations for any feature in our dataset are possible.

#### Predicting DOC and DTN with hybrid SL models

Hybrid microbiome-biogeochemistry datasets make more accurate predictions than biogeochemistry or the microbiome, alone. We use the RMSE of residuals in addition to simple predicted value vs. observed value scatter plots to evaluate the accuracy of statistical models. Here, we report median log10-space testing data RMSEs of 0.23 mg l^-1^ and 0.22 mg l^-1^ for dissolved organic carbon (DOC) and dissolved total nitrogen (DTN), respectively, from a hybrid biogeochemistry and microbiome feature space (Table 1). Given the broad range of DOC and DTN observed in this study (Supplementary Table 5), these accuracies are notable. We mitigated the effect of outlier predictions on model RMSEs through cross-validation and report the median performance of 10 different training / testing data splits (Table 1). We also visualized prediction vs. observation scatter plots to facilitate intuitive assessment of model accuracy (Fig. 2 & Fig. 3). Cross-validation determined which biogeochemical and microbiological features are important for making predictions (Fig. 4). Examination of coefficients selected during SL shows that, for hybrid dataset models, fewer biogeochemical terms are used than in biogeochemistry-only models; microbiome data are favored and assist in higher accuracy predictions of biogeochemical responses. Further, selection of useful microbiological features for making predictions shows that the rare biosphere is more useful than the abundant biosphere (Fig. 5).

**Fig. 4.**
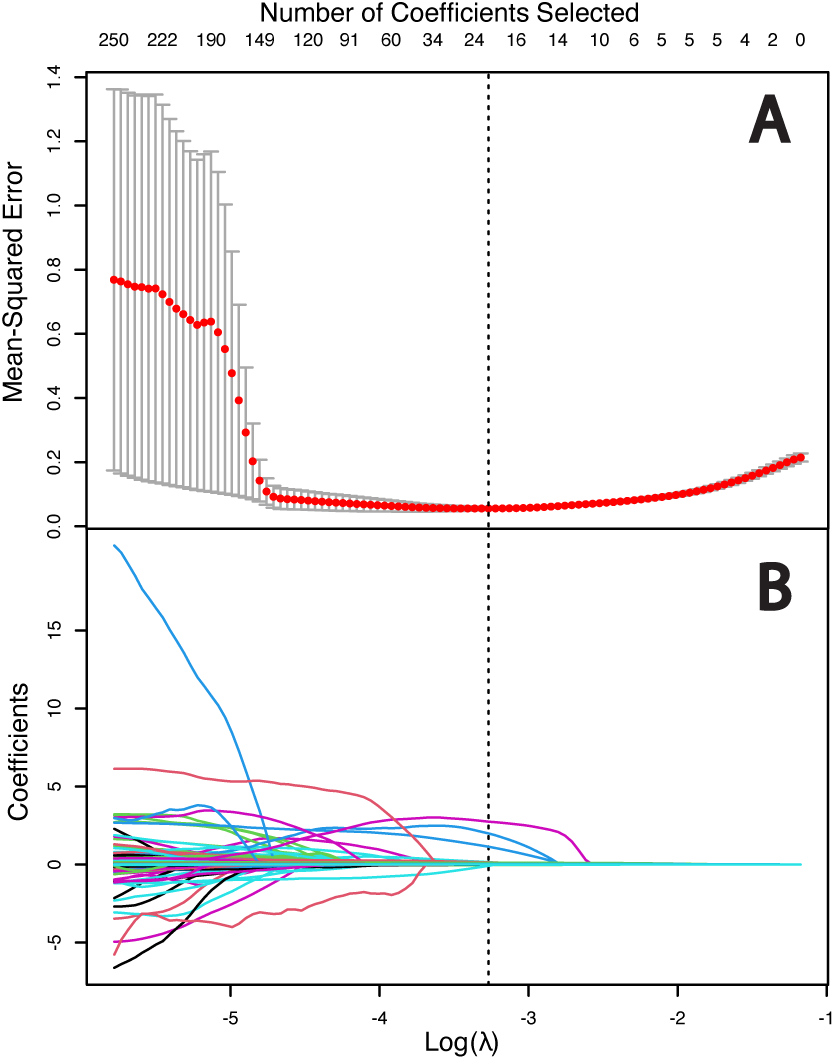
Statistical learning (via lasso-penalized linear regression) of the best features—and their coefficients—for predicting leachable dissolved total nitrogen (DTN) from soil after wildfire with a hybrid SL model. The objective was to learn a useful set of parameters while discarding unhelpful ones; both too many and too few features included in the final model result in an undesirable accuracy. (A) Learning the optimal penalty weight (*λ* on x-axis) that coefficients must overcome to enter the model (dotted line) (Equation 1); this is done by identifying the *λ* that minimizes the mean model error (red points) with 10-fold cross validation (gray error bars are +/- standard error). (B) There is an optimal number of selected coefficients (each colored line represents a feature) in the hybrid model that produces the best model fit (dotted line), and this combination is an ensemble of biogeochemical and genera microbiome terms. The coefficients of features shrink to *exactly* 0 (unimportant) with increasing *λ*. The optimal value of *λ* (dotted line) is large enough to remove noise from the dataset—overcoming the statistical nuisance of variance inflation—but small enough to retain enough complexity necessary to make accurate predictions.

**Fig. 5.**
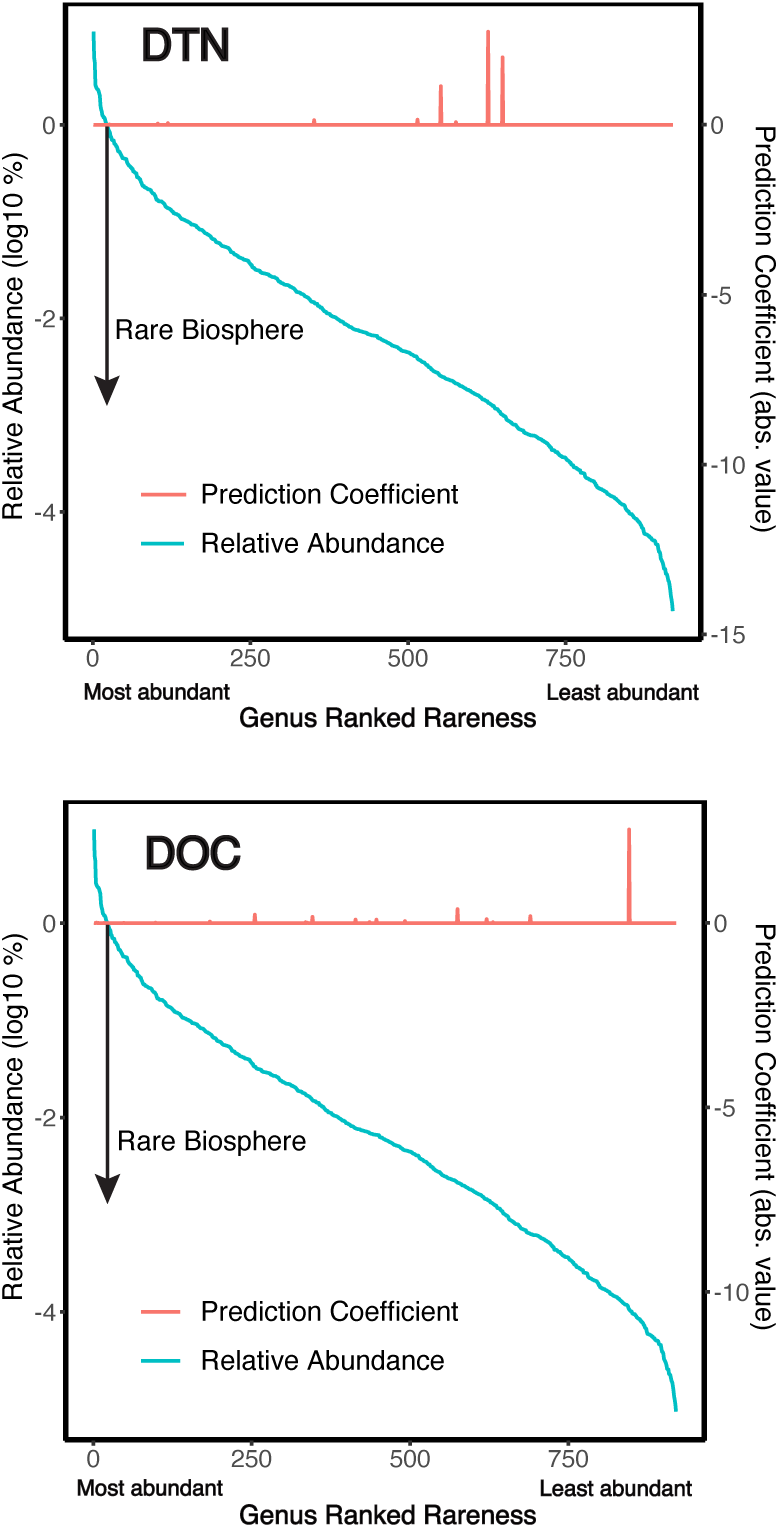
The contributions of rare genera of soil microorganisms to effective hybrid model predictions of water-extractable dissolved total nitrogen (DTN) and dissolved organic carbon (DOC). With decreasing abundance of genera of microorganisms (blue curve), their predictive utility, as measured by their coefficients in the learned linear model (red peaks), becomes greater. A prediction coefficient value of zero means that the microorganism was not included in the learned model at all. The black arrow begins at the threshold for rare microbiota on the left y-axis (relative abundance) and indicates the direction of increasing rareness—any organism with a log10 % relative abundance of *<* 0 is rare (*<* 1% mean relative abundance across all samples in the dataset). The vast majority of non-zero coefficients (important features) come from the rare biosphere, though some abundant microbiota still have extremely small prediction coefficients on this figure.

#### Predicting NH_4_^+^ with only the microbiome

To further probe whether or not the the soil microbiome, alone, is useful for explaining biogeochemical responses (with no biogeochemical or SBS information), we conducted a case study of NH_4_^+^ —a biogeochemical analyte that typically increases after wild-fire but is quickly depleted in the following months and years (Supplementary Tables 2 & 4) [52–54]. First, we tested for the effect of SBS on NH_4_^+^ concentrations across all soil samples (one-way ANOVA: p = 1.67 *×* 10^−4^). Since NH_4_^+^ varies significantly by SBS (an important general measure of fire impact on soil), we explicitly tested whether or not the median NH_4_^+^ concentration for each SBS class is predictive of biogeochemical response (Fig. 6).

**Fig. 6.**
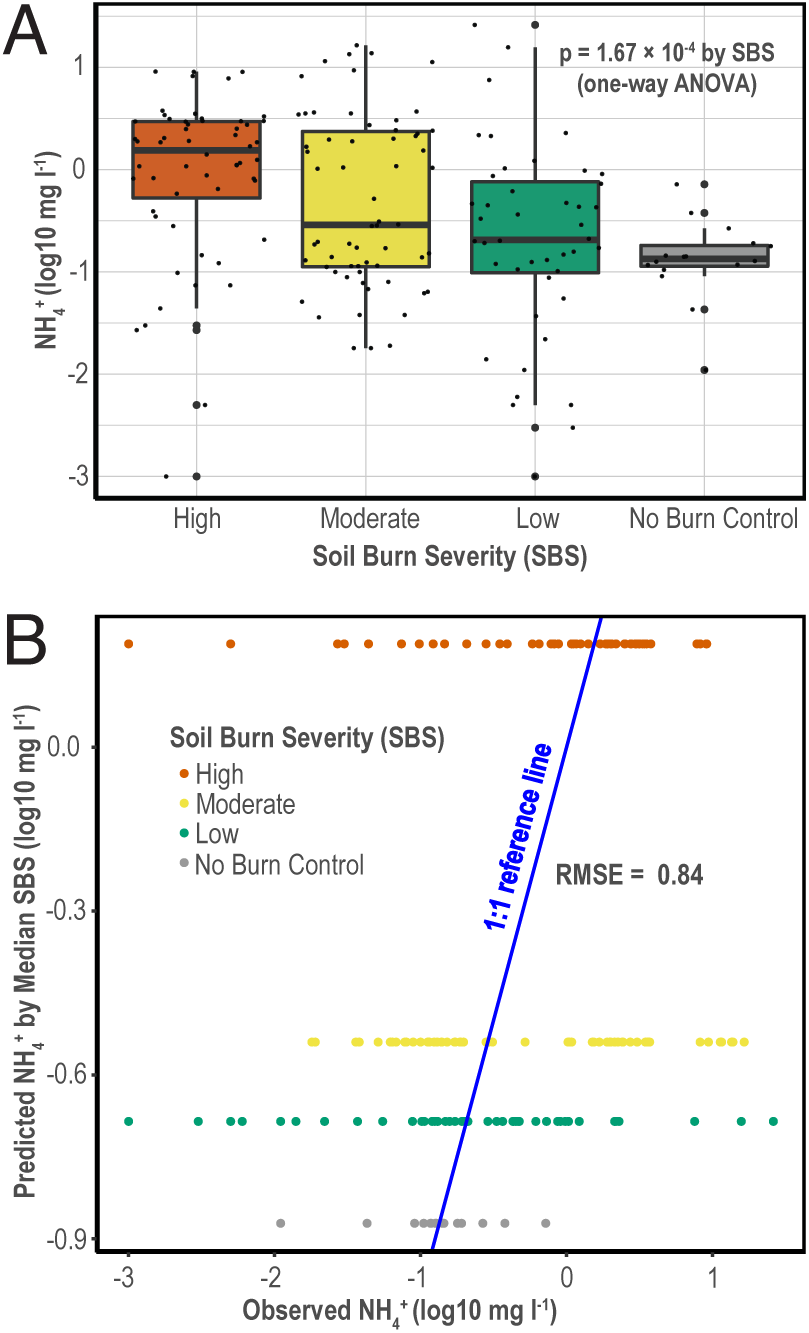
Although NH_4_^+^ varies significantly by SBS, using SBS to make explcit predictions about NH_4_^+^ response results in poor accuracy. (A) A box and whisker plot of NH_4_^+^ concentration for each SBS; these variations are significant (one-way ANOVA: p = 1.67 × 10^−4^). (B) Using the median NH_4_^+^ by SBS to predict NH_4_^+^ response. In a successful prediction model, data points would follow the 1:1 prediction line. Although, on average, there is significant variation in NH_4_^+^ that is explained by SBS (panel A), using SBS alone to predict NH_4_^+^ response results in poor accuracy (as shown by horizontal scatter in panel B). SBS is important in a descriptive sense to understanding generalized biogeochemical responses after wildfire, but a multi-dimensional statistical model is required to accurately predict explicit NH_4_^+^ concentrations.

Despite the significant differences in NH_4_^+^ among SBS classes, SBS has poor accuracy as a predictor of NH_4_^+^ concentration. This finding is unsurprising as there are only four discrete levels of burn severity from which to make predictions (High, Moderate, Low, and No Burn); nonetheless, it explicitly motivates the need for SL models that exploit nuance to make more accurate predictions. Next, since elevated NH_4_^+^ post-fire (relative to unburned soil, Supplementary Tables 2 & 4) provides an energetic nitrogen source that chemolithoautotrophic ammonia-oxidizing-Archaea (AOA) [55–59] and ammonia-oxidizing-Bacteria (AOB) [60–62] may exploit—and this is supported by concomitant elevations in nitrate (NO_3_^-^) in the same soils—we examined the relationship between NH_4_^+^ concentration and AOA/AOB [63]. To test this relationship, we regressed NH_4_^+^ on AOA/AOB relative abundances and found no correlation (Fig. 7). Further, elevated NH_4_^+^ and NO_3_^-^ concentrations in burned versus unburned soils immediately post-fire are rapidly depleted in the first and second post-fire years (44% and 74% losses in NH_4_^+^ and NO_3_^-^, respectively, in high SBS Org-horizons at the 416 Fire in the first year) which suggest that impacts on nitrogen nutrients from geophysical / hydrological processes are substantial. Neither SBS nor AOA/AOB, alone, explain NH_4_^+^ response immediately post-fire and in the post-fire years; far more information is likely necessary.

**Fig. 7.**
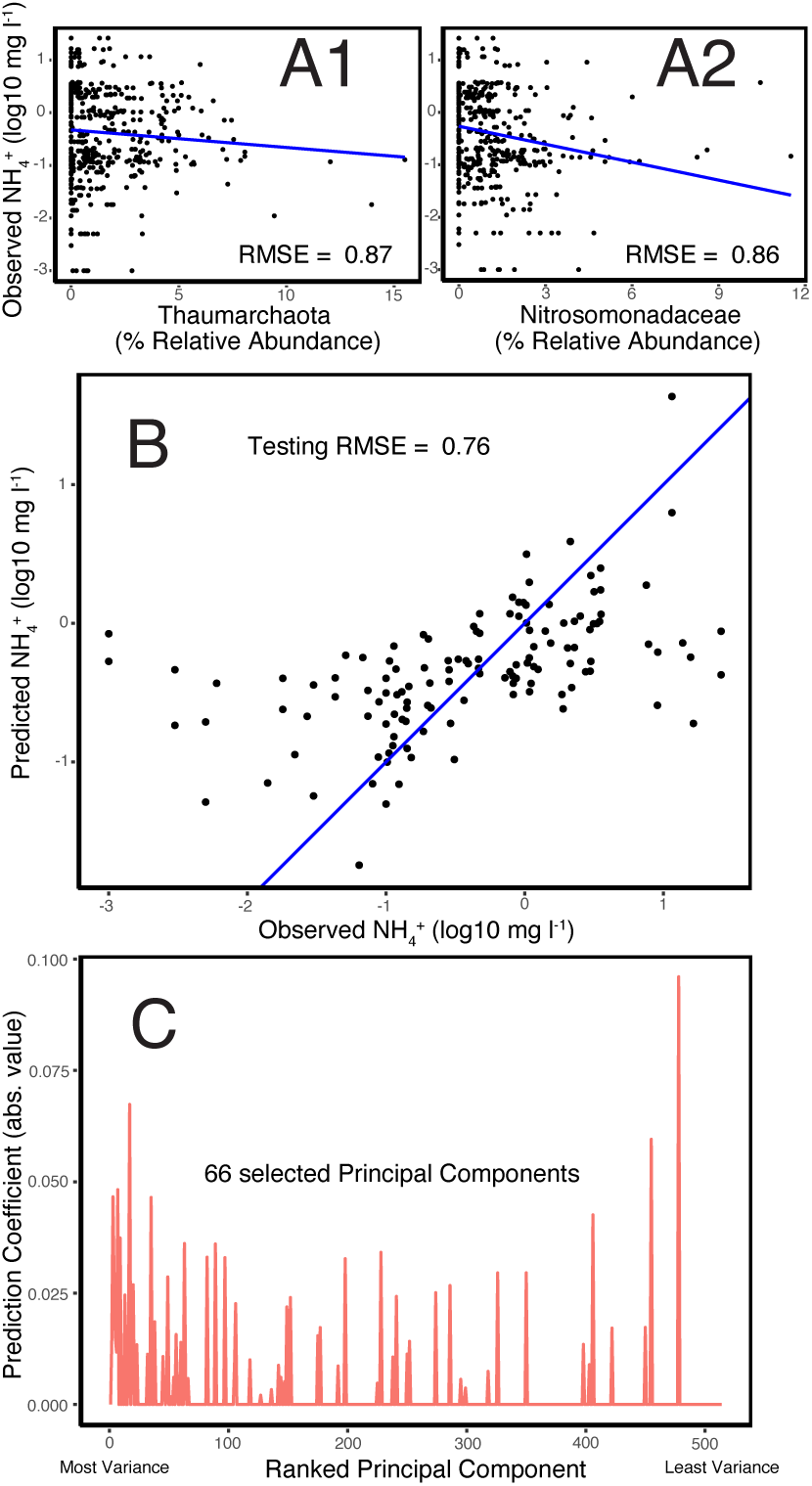
Predicting and explaining soil ammonium (NH_4_^+^) in the post-fire years using only the microbiome. (A1) There is no relationship between NH_4_^+^ and the abundance of canonical ammonia-oxidizing-Archaea (AOA) from phylum Thaumarchaeota in soil (a poor linear fit, blue line); 100% of the Thaumarchaeota sequenced in our study were of class Nitrososphaera. (A2) The linear fit (blue line) between ammonia-oxidizing-Bacteria (AOB) from family Nitrosomonadaceae and NH_4_^+^ is also poor. (B) However, NH_4_^+^ response is well-explained by PCA regression. The blue line is a 1:1 reference for predicted (using the PC regression model) vs. observed NH_4_^+^. The model was trained on 75% of samples, and the performance of the model on test samples is plotted here (the remaining 25% of data). The spread of data about blue lines is critical to assessing model accuracy; RMSE reports the error of the model in log10-space and so small differences in RMSE are substantial differences in the accuracies of models. (C) Surprisingly, many PCs were selected during PCA regression, suggesting that microbiome complexities (and not just abundant microbiota) are critical to accurately predicting NH_4_^+^ response in the post-fire years. NH_4_^+^ can be explained by an integrative approach to unpacking the microbiome; the presence of the usual suspects of NH_4_^+^ oxidation (AOA/AOB) do not explain the response of NH_4_^+^ with the same skill as SL. PC regression overestimates NH_4_^+^ at low concentrations, and tends to underestimate it at high concentrations. Low concentrations may be hard to predict in general due to instrument detection limits, but high concentrations might be better predicted with the development of more nuanced statistical models that allow for non-linear biogeochemical relationships.

Finally, we tested whether or not the microbiome alone and SL accurately predicts NH_4_^+^ response (Fig. 7). The microbiome was examined by extending our lasso-penalized linear statistical model by first conducting principal component analysis (PCA) on the data. PCA is a dimension reduction technique that learns a relatively small set of *new* features (principal components —’PCs’) that are linear combinations of all features in the dataset. If only a few PCs are necessary to explain data, then the system is not complicated. We regressed NH_4_^+^ on PCs with lasso-penalized linear regression (trained on 75% of samples and tested on the remaining 25%) to determine how many PCs are important. Surprisingly, dozens of PCs were selected and predictions on testing data were accurate —indicating that considering only the dominant microbiota in explaining responses is insufficient. Paradoxically, although the microbiome can predict enriched NH_4_^+^ concentrations in soil post-fire, this process is not thought to be biological; NH_4_^+^ derives from combusted vegetation and Org-horizons [52–54]. However, in this case study of SL with principal component regression, we found that the microbiome provides the sufficient complexity required to predict NH_4_^+^. This suggests that even abiogenic analyte pools such as NH_4_^+^ after wildfire can be explained by microbial tenants [64] of an ecosystem —independent of whether or not those microbiota actually produce or consume the analyte. These tenants may be responding to other environmental changes post-fire (e.g., increases in pH and acid-neutralizing capacity (ANC)), but their aggregate responses to these changes still aid in the prediction of NH_4_^+^ through a complex web of co-variances and inter-dependencies. It is possible to use the microbiome to predict NH_4_^+^ concentrations both immediately post-fire and in the post-fire years, and this SL approach provides far more accurate predictions than the descriptive use of either SBS or microbiota that have been shown to actively oxidize NH_4_^+^ (AOA/AOB) in soils [63].

## 5 Discussion

Soils are critical to understand due to their essential roles in maintaining ecosystem productivity [65, 66]. Fire is a natural part of many ecosystems, but it is unclear how soils and their microbiomes will respond to shorter fire return intervals and combinations of disturbances and climatic stress in burned landscapes [1, 3, 67]. Here, we conducted a study that examines a large multivariate dataset from wildfire-impacted soil and show that biogeochemical responses can be explained with linear SL models. To our knowledge, there are no datasets of similar construction on which we could further test and validate our models; however, we have extensively evaluated predictions on held-out testing data from two different fires/forests.

We learned what aspects of the soil biogeochemical world are important to explaining the variance in soil at field scale. We find that hybrid microbiome + biogeochemical models are the most accurate (Table 1). Further, most microbial features selected by SL as useful for making accurate predictions of biogeochemical responses are uncommon (*<* 1% relative abundance at any phylogenetic rank) (Fig. 5 and Supplementary Tables 7 - 10). We can use SL to exploit complexities to make predictions about specific soil responses in this burned environment. There is likely a far greater density of information in soil biogeochemistry and the soil microbiome that can be leveraged for applied inference and prediction than previously recognized.

For particularly important ecosystem features such as NH_4_^+^ [4, 54] it is worth comparing the predictive power of physical descriptions (such as SBS) as well as NH_4_^+^-associated microbiota (AOA/AOB) to the accuracy of SL models. Although NH_4_^+^ varies significantly as a function of SBS, we find that SBS is not *predictive* of NH_4_^+^ response (Fig. 6). Members of the AOA phylum Thaumarchaeota and AOB family Nitrosomonadaceae are common in our dataset (Figs. 1 & 7) and their ability to use ammonia for chemolithotrophy is important to global nitrogen cycling in both marine and terrestrial systems [55–63, 68–70]. Nonetheless, AOA/AOB relative abundances do not *predict* NH_4_^+^ response. The presence of AOA and AOB in the soils we analyzed is a case in point of the potential of SL in understanding the biogeochemistry of a *system* (Fig. 7).

Environmental scientists acknowledge that environmental systems are enormously complex, yet we often aim to distill that complexity to a handful of what we think are principal, controlling features. This dissonance is especially acute as the scale of the system increases. One might expect a correlation, even if weak, between the relative abundances of AOA/AOB and NH_4_^+^ since these microbiota can exploit NH_4_^+^ for energy and/or nitrogen [71], but we do not observe this relationship (Fig. 7). Though genomes/transcriptomes of AOA/AOB in our study would be necessary to fully explain their activity, a possible alternative view supported by our SL analysis is that cost-efficient, high-throughput microbial community profiling with the SSU rRNA gene may be sufficient if the goal is strictly to predict responses in soil (a large *N* from high throughput sequencing may compensate for the loss in large *p* offered by multi-omic methods). Gene-specific investigations of post-fire soils found that neither *nifH* nor *amoA* (the genes that encode the enzymes for nitrogen-fixation and ammonia-oxidation, respectively) were consistently expressed or even present [5, 72, 73]. A functional model of post-fire soil may not predict DTN or NH_4_^+^ responses if the enzymes responsible for their generation/transformation are not consistently quantifiable—yet DTN and NH_4_^+^ post-fire vary greatly on a continuous scale (Supplementary Tables 1 - 5).

As one possible solution, we show it is feasible to leverage the composition of the total microbiome cost-efficiently with hundreds of SSU rRNA gene ampilcons to explain DTN and NH_4_^+^ (and other) responses in the first and second post-fire years. A critical distinction is that this does not mean that the microbial community is necessarily responsible for generating or consuming NH_4_^+^ post-fire; in fact, elevated NH_4_^+^ is a typical product of organic matter oxidizing in wildfire [52, 53] and post-fire erosion and plant-uptake are additional sinks for NH_4_^+^ [74, 75]. We show that independent of how NH_4_^+^ is mechanistically regulated in post-fire soil, it is possible to use the impressive data density of DNA to predict responses in the first two post-fire years without microbial genomes or transcriptomes (though multi-omic methods are certainly valuable and *necessary* to a detailed molecular description of environmental processes). The regulation of NH_4_^+^ and other analytes post-fire is likely dependent, in ways still unknown, on the complex milieu of geophysical processes and macroscopic organisms as well as competition, symbioses, and horizontal gene transfer in the microbial world —even if redox reactions themselves are carried out by a small set of core genes [76].

The power of SL, coupled with an appropriately-large dataset, is the ability to explore complex systems in a heuristic fashion to possibly discern patterns and relationships that were previously unknown or unknowable in the absence of the SL techniques. SL is unaware of the meanginfulness of features to domain scientists —and therein lies an opportunity to generate new hypotheses and discover unexpected phenomena (such as the predictive power of the microbial rare biosphere; Fig. 5). Features possibly unrelated to microbial activity immediately after fire (e.g., NH_4_^+^) can still be explained with surprising accuracy by quantifying the response of the microbial community—even if these responses are primarily governed by other harsh fire/post-fire conditions (e.g., excessive heat and increases in soil pH). We hypothesize that the composition of microbial communities in soil records the life-history and potential of an environment —and this record is valuable for making predictions about innumerable biogeochemical responses.

The rare biosphere is critical to making successful predictions (Table 1 & Fig. 7) while SBS and AOA/AOB relative abundances, by themselves, are unhelpful (Fig. 6). The microbial rare biosphere (such as that we characterized shortly after wildfire) is poorly understood—yet likely serves as a nearly inexhaustible source of genomic innovation as described by Sogin and colleagues in 2006 [77]. We suggest that the rare biosphere, and the inherent complexities of diverse microbial communities in general, possess the information necessary for accurate biogeochemical predictions and explanations as learned by statistical models (Table 1). Microbial communities and their diversity/traits are important to the resilience and trajectory of ecosystems [78, 79], but quantifying these attributes to explain variation in soil is an ongoing challenge [80]. A new approach to describing complex systems may be necessary in environmental science—one that exploits, rather than constrains, the variability of natural systems. Such an approach may yield true predictive capability in the environment while simultaneously elucidating impactful, unexpected, mechanisms to test experimentally. The feasibility of explicit, accurate, predictions for soil responses suggests that the potential, trajectory, and success of post-fire restoration interventions could be quantitatively evaluated in an SL framework —possibly enabling the optimization of restoration strategies for impact and cost-effectiveness in environmental policy. Additional research should investigate the feasibility of using microbiome and biogeochemical data to forecast future responses in environmental systems.

Leveraging the complexities of environmental systems as an asset [81] with SL, ML—or Artificial Intelligence (AI), in general—likely has numerous practical applications. As with post-wildfire soils, agricultural soils [82], biological wastewater treatment systems [83], and oceans [84] all have critical biogeochemical cycles that require greater understanding. Global microbial-biogeochemical systems [76, 85] offer a near inexhaustible supply of data density—possibly enabling the principles of biomedical ’personalized medicine’ to be extended to environmental science for the precision management of soil, water, and air.

## Data and Code Availability

Our data are FAIR (Findable, Accessible, Interpretable, and Reusable) [86]. All raw DNA reads are deposited at the NCBI SRA under BioProject PRJNA767436. All biogeochemical data, post-processed DNA sequence data, metadata, and code are available on Zenodo (DOI: 10.5281/zenodo.5539890). During review, the Zenodo DOI is reserved and NCBI data are not public. Our FAIR data and code are sufficient to reproduce all of our analyses. Our code is broken down into one preprocessing file, one analysis file, and one file used to make Earth maps—all as RMarkdown files (.Rmd) with comments and explanations for every step. SL modeling of any analyte in our study can be done within our analysis code, and figures analogous to what we present in this manuscript can be automatically produced. Interested parties can explore their own questions relevant to our dataset. Our full analysis pipeline, including figure generation, is reproducible in one hands-off run of code in *∼* 1 hr on a 2018 MacBook Pro.

## Supplementary Information

This research article has an accompanying Supplementary Information.

## Author Contributions

Conceptualization: A.S.H. Methodology: A.S.H. and J.R.S. Field Investigation: A.S.H., H.F.P., and J.R.S. Biogeochemistry Investigation: T.S.F., A.S.H., and C.C.R. Molecular Biology Investigation: A.S.H., H.F.P., N.A.M., and J.R.S. Software: A.S.H. Formal Analysis: A.S.H., D.C.V., and W.K. Resources: J.R.S. and C.C.R. Writing - Original Draft: A.S.H. Writing - Review and Editing: All authors. Supervision: J.R.S. Funding Acquisition: A.S.H. and J.R.S.

## Acknowledgments

We are grateful to Dr. Laura Leonard and Dr. Michael Remke for assistance with field work, to Dr. Blake Stamps for molecular biology technical advice, to Dr. James Ranville and Shaun Bevers for helpful discussions about geochemistry, and to Kalen Rasmussen and Michael Vega for assistance with DNA sequencing preparation. Henry and Lin Ballard, Eric Bader, and Chief Michael Schmitt (Sunshine Fire, Boulder County, Colorado, USA), Maya MacHamer (Fourmile Watershed Coalition, Boulder County, Colorado, USA), and Jared Whitmer (Columbine Ranger District, San Juan National Forest, USA) generously lent their time to help us identify and gain access to active wildfire. The Columbine Ranger District of the San Juan National Forest and the Salida Ranger District of the San Isabel National Forest graciously permitted our access to burn areas. This research was supported by the National Science Foundation Graduate Research Fellowship Program (to A.S.H., Fellow I.D. #2019258966), and NASA Exobiology research grant #80NSSC19K0479 (to J.R.S.).

