## Supplementary Information for "Statistical learning and uncommon soil microbiota explain biogeochemical responses after wildfire"

### 1 Supplementary Wildfire Sampling Methods

#### 1.1 416 Fire Field Methods

Preliminary Burn Area Emergency Response (BAER) Team and burn area reflectance classification (BARC) maps were used to navigate to a region of the fire that: 1) represented a transect that contained all SBS classes (No Burn Control, Low, Moderate, and High); and 2) was not involved in ongoing firefighting operations. 17 unique plots were established and on-site SBS was determined by examining the extent of burned vegetation and combustion in the Org and Min horizons at each plot [1]. No Burn Control plots were selected from unburned areas adjacent to burned plots by attempting to match pre-fire vegetation; however, pre-fire vegetation guides where fire burns and so representative control soils are challenging to define. Individual plots along SBS transects were selected by avoiding vegetation and balancing the number of samples for each SBS class; aspect, slope, and pre-fire vegetation type were not constrained. The distance between plots was maximal given the constraints of vegetation, balancing sample numbers by SBS class, and active wildfire. 30 cm pits were dug at each sampling location. One Org-horizon and multiple, depth-sequential/evenly spaced, Min-horizon samples were scooped into clean 590 mL Ziploc™ plastic containers, sampling horizons from bottom-up to avoid cross-contamination. Bulk soil was thoroughly homogenized with a clean scoopula prior to biological triplicate collection for DNA (with re-homogenization of bulk soil between samplings). Remaining bulk soil (singlicate) was saved for biogeochemistry.

#### 1.2 Decker Fire Field Methods

Only Org-horizon samples were collected. 30 unique plots across a SBS gradient were selected with the same plot selection and soil sampling methodology as for the 416 Fire.

#### 1.3 Sample Collection and Storage Specifics

*DNA Samples:* Triplicate 0.25 g samples of 2 mm size-fraction homogenized soil were collected into either 2 mL cryogenic storage vials or ZR BashingBead™ Lysis Tubes from the ZymoBIOMICS™ DNA Miniprep Kit that were preloaded with 750 µL of Zymo DNA/RNA Shield™ to preserve nucleic acids. All microbiome soil collection vials were stored on dry ice in the field for a maximum of 48 hours before definitive storage in the lab. In lab, samples in DNA/RNA Shield™ were stored at room temperature per manufacturer instructions and samples in cryogenic storage vials were frozen at -80° C.

*Biogeochemistry Samples:* After field triplicate microbiome sample collection, a 100 mL aliquot was subsampled from the homogenized bulk soil into 50 mL Falcon conical centrifuge tubes (Thermo Fisher Scientific, Waltham, MA, USA) for subsequent biogeochemical analysis. Bulk soil aliquots were kept on ice for a maximum of 48 hours and then stored at 4° C in the lab.

### 2 Supplementary Biogeochemistry Methods

#### 2.1 Analytical Characterization of Filtrates

Anion and cation concentrations were determined by ion chromatography with electrolytic suppression and conductivity detection with an AG19A guard column and an AS19A ion-exchange column for anions, and a CG12A guard column and a CS12A ion-exchange columns for cations (Thermo-Fisher Corporation, Waltham, MA). Detection limits were 0.01 mg $l^{-1}$ for $Ca^{2+}$, $Cl^{-}$, $K^{+}$, $Mg^{2+}$, $Na^{+}$, $NH_4{}^{+}$, $NO_3{}^{-}$, $SO_4{}^{2-}$, and $PO_4{}^{3-}$. Acid neutralizing capacity (ANC) was measured on unfiltered subsamples by Gran titration (Gran 1952). Electrical conductance and pH were analyzed automatically with Mettler Toledo sensors (Mettler Toledo Corporation, Columbus, OH). Dissolved organic carbon (DOC) and Dissolved total nitrogen (DTN) were determined using a Shimadzu TOC-$V_{CPN}$ total organic carbon analyzer, with 2 M HCl addition before analysis to remove mineral C (Shimadzu Corporation, Columbia, MD). Detection limits for DOC and DTN were 0.1 mg $l^{-1}$. Dissolved organic nitrogen (DON) was calculated by subtracting the sum of $NO_3{}^{-}$-N and $NH_4{}^{+}$-N from DTN.

### 3 Supplementary DNA Sequencing Methods

#### 3.1 Reaction Concentrations and Primers

Per 50 µL PCR reaction: 1x QuantaBio 5Prime HotMasterMix, 0.2 µM 515Y-M13 forward primer (5′-GTA AAA CGA CGG CCA GT CCG TGY CAG CMG CCG CGG TAA-3′), 0.2 µM 926R reverse primer (5′-CCG YCA ATT YMT TTR AGT TT-3′), and template genomic DNA (gDNA). The underlined portion of the forward primer is a linker sequenced used to attach unique barcodes during an additional 6-cycles of PCR for DNA sequencing [2–4]. Due to concentrated organic acids in wildfire samples [5] PCR amplification was optimized through serial dilution series of extracted input genomic DNA (gDNA) at 1:10, 1:100, and 1:1000 concentrations; on a per-sample basis, the dilution series that resulted in the best yield of PCR product was used in downstream steps. PCR parameters were as follows: 1 cycle of initial denaturation at 94° C for 2 minutes; 30 cycles of denaturation at 94° C for 45 seconds, annealing at 50° C for 45 seconds, and extension at 68° C for 1 minute 30 seconds; 1 cycle of a final extension at 68° C for 5 minutes; a final hold at 4° C. The 6-cycle PCR barcoding step had the same reaction concentrations and thermocycling conditions as the initial PCR. PCR products from both initial amplification and barcoding were cleaned and size-selected to the target amplicon size with a 0.8× volume of KAPA Pure Beads (Roche Sequencing, Pleasanton, CA).

### 4 Supplementary dPCR Methods

#### 4.1 Primers

Bacteria, Archaea, and microbial Eukarya absolute quantification was determined by domain-specific primers for each. *Bacteria*: Eub338 (5'-ACT CCT ACG GGA GGC AGC AG-3') and Eub518 (5'-ATT ACC GCG GCT GCT GG-3'); 53° C annealing temperature [6]. *Archaea*: Arch967F (5'-AAT TGG CGG GGG AGC AC-3') and Arch1060R (5'-GGC CAT GCA CCW CCT CTC-3'); 50° C annealing temperature for 30 seconds [7]. *Microbial Eukarya*: 5.8s (5'-CGC TGC GTT CTT CAT CG-3') and ITS1f (5'-TCC GTA GGT GAA CCT GCG G-3'); 53° C annealing temperature [6].

#### 4.2 Reactions, Thermocycling, and Imaging / Quantification

DNA samples were diluted 1:10 prior to dPCR due to over-fluorescence in non-diluted samples. dPCR has been shown to outperform quantitative PCR (qPCR) in the absolute quantification of microbial communities in environmental samples that can have low biomass, contain PCR-inhibiting organics, or both [8, 9]. Empirically, we found some wildfire samples can be low biomass; further, most samples likely contain at least some PCR-inhibiting humic/fulvic acids [5] which would explain the dilution series necessary to get many samples to amplify during amplicon PCR for DNA sequencing.

dPCR reactions for all three domains of life followed manufacturer recommendations for the QIAcuity EG PCR Kit, and these were: 4 µL 3× Master Mix, 2 µL 1:10 diluted template gDNA, 0.4 µM forward primer, 0.4µM reverse primer, and PCR water to a final reaction volume of 12 uL. *Bacteria Thermocycling*: 1 cycle of 95° C for 2 minutes; 30 cycles of 95° C for 30 seconds, 53° C for 30 seconds, and 72° C for 45 seconds; 1 cycle of 40° C for 5 minutes. *Archaea Thermocycling*: 1 cycle of 95° C for 2 minutes; 30 cycles of 95° C for 30 seconds, 50° C for 30 seconds, and 72° C for 45 seconds; 1 cycle of 40° C for 5 minutes. *Microbial Eukarya Thermocycling*: 1 cycle of 95° C for 2 minutes; 30 cycles of 95° C for 30 seconds, 53° C for 30 seconds, and 72° C for 1 minute 15 seconds; 1 cycle of 40° C for 5 minutes. The same imaging parameters were used for all three domains of life: Green channel, 300 ms exposure, gain of 6. The threshold relative fluorescent units (RFUs) were chosen on a plate-by-plate basis according to the following manual protocol: The sample on the plate with the negative detection population that had the highest RFUs was used to pick the threshold. The threshold was set directly above the point cloud of negative wells on this sample, and then applied universally across all samples on the plate. This is a conservative thresholding choice, meaning that across all micro-wells for any given sample on the plate, there must be a high degree of confidence that the RFUs are bright enough before any given micro-well is classified as a 'positive'. The thresholding values (RFUs) for each plate were: 49.24 for Bacteria, 24.88 for Archaea, and 49.55 for Eukarya. Since dPCR does not use a standard curve to calibrate observed counts of gene copies, on a per-plate basis we calculated the mean number of counts observed across all negative controls and subtracted that number from the measured counts in true samples.

### 5 Supplementary Bioinformatics Methods

#### 5.1 Sequence Data Pre-Processing

Raw paired-end DNA sequence reads were demultiplexed with Adapter Removal v2 [10]. Primers were removed from reads, paired-end reads were merged (Bacteria/Archaea independently from Eukarya [4]), and sequence variants were inferred for individual sequencing runs with Dada2 [11]. Learned amplicon sequence variants (ASVs) from independent sequencing runs were combined into one table (for each of Bacteria/Archaea and Eukarya) for chimera removal and taxonomy assignment (Bacteria/Archaea: Silva nr99 v138. Eukarya: Silva nr v132) by Dada2 [11–13]. Class Nitrososphaera is classified under phylum Crenarchaeota in Silva nr99 v138; however, updated phylogeny classifies Nitrososphaera as being from phylum Thaumarchaeota [14, 15]. Thus, we manually assigned all Nitrososphaera ASVs to phylum Thaumarchaeota in figures. ASVs, taxonomic calls, and sample metadata were combined into a single data structure object with Phyloseq [16]. Bacteria/Archaea and Eukarya phyloseq objects serve as the base input file for all downstream code and analyses.

#### 5.2 Preparation for SL

The Bacteria/Archaea and Eukarya phyloseq objects were merged into a single dataset for simultaneous consideration of all three domains of life. Chloroplast and Mitochondria sequences were removed. Arthropoda (and other non-microbial Eukarya) DNA sequences were retained in the dataset since they represent real information from soil. However, these few non-microbial reads have small mean relative abundances; for example, Arthropoda sequences have a mean relative abundance of only 0.08 %. The total DNA sequence dataset is comprised of 8 209 175 sequences (222 374 of which are microbial and non-microbial Eukarya, combined). Contaminating reads were identified and removed from samples on a per-sequencing-run basis with Decontam [17]; choosing either contaminants or *not* contaminants as the null hypothesis was determined by *a priori* knowledge of the biomass of samples, and decision thresholds were chosen by examining a histogram of model scores for each sequencing run, as recommended by the Decontam documentation. Relative abundances of taxa at each phylogenetic rank were computed with ampvis2 [18].

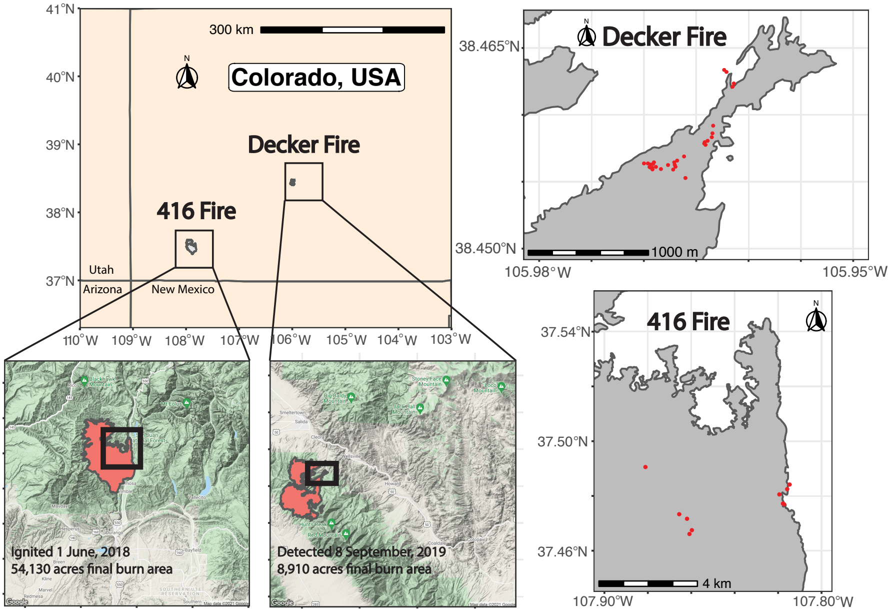


**Supplementary Fig. 1** Wildfire study sites. Left: An overview of southwest Colorado, USA and, in red, the burned area polygons of the 416 and Decker fires. Right: Map detail of sampled regions within each burn area; red points are recurrent sampling locations.

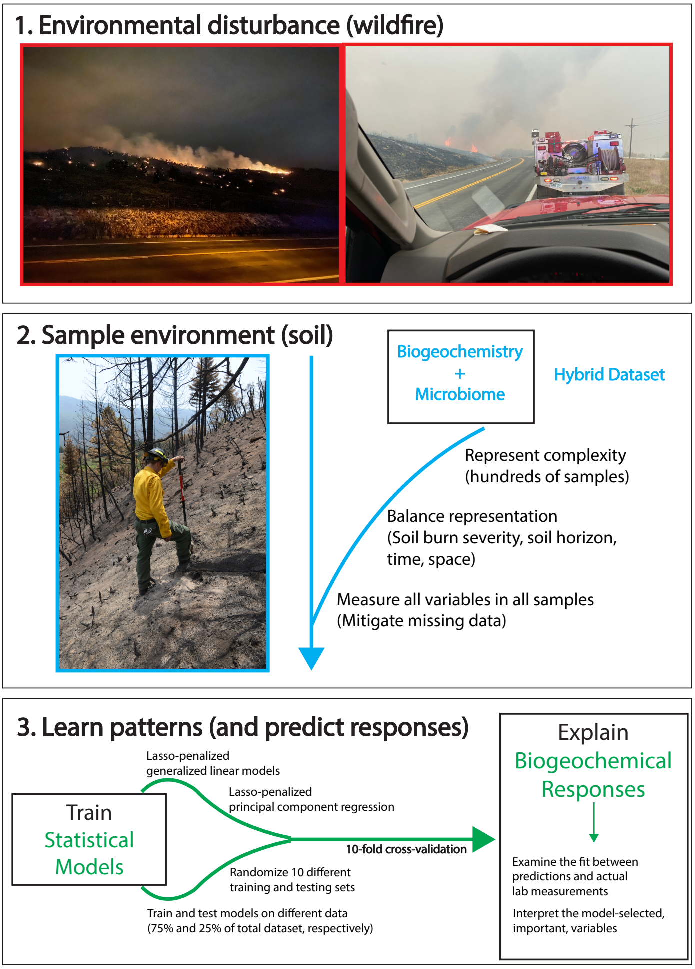


**Supplementary Fig. 2** Conceptual process-flow for the study. (1) Wildfire targeted for sampling within weeks after active burning. (2) Soil samples were taken for both biogeochemistry and microbiome quantification in the lab; critical environmental factors were thoroughly represented in samples. (3) Statistical models were trained and cross-validated on 75% of the data to learn patterns associated with biogeochemical responses. Learned models were tested on the remaining 25% of held-out data to evaluate accuracy.

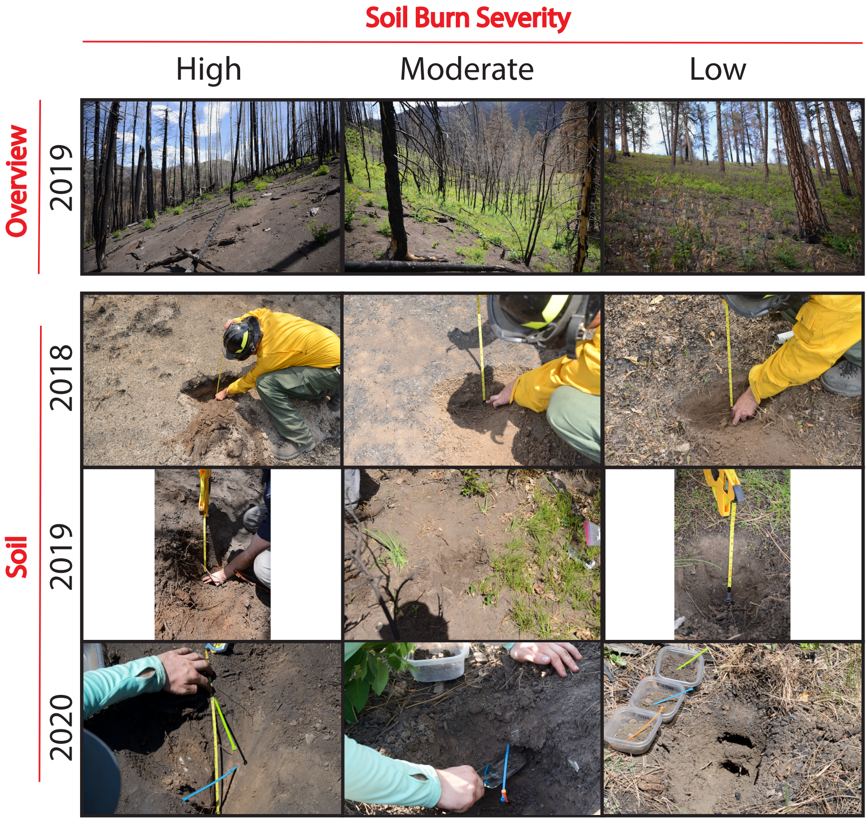


**Supplementary Fig. 3** Representative overview plots and soil depth profiles from the 416 Fire, Durango, Colorado. One plot of each soil burn severity (SBS) is depicted over 3 summers. There is substantial regrowth of vegetation at the Moderate and Low SBS sites in the years following wildfire, though some recovery is seen at High severity sites as well.

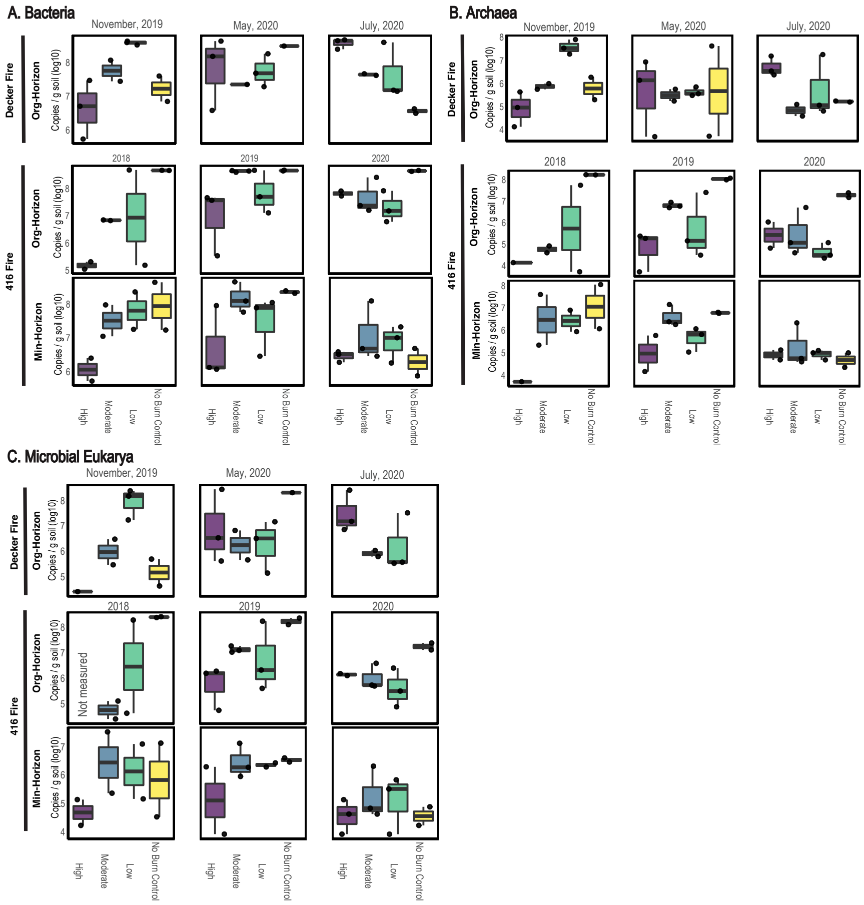


**Supplementary Fig. 4** Absolute quantification (via dPCR) of microbial marker gene copy numbers in soil across all three domains of life post-fire. Reported numbers are copies of marker gene per gram of soil. The box and whisker plots depict median, 1$^{st}$ and 3$^{rd}$ quartiles, and the largest and smallest values. The number of marker genes in soil is different by soil burn severity (SBS; as High, Moderate, Low, and No Burn) and organic versus mineral horizon (Org/Min-Horizon). Highly burned soil is not sterile. dPCR is known to better quantify microbial load in low biomass samples with PCR inhibitors [8, 9]; our detection of genes here supports the legitimacy of non-contaminating reads that were sequenced. (A) Universal Bacteria primers. (B) Universal Archaea primers. (C) Microbial Eukarya primers. See Supplementary dPCR Methods for primer sequences.

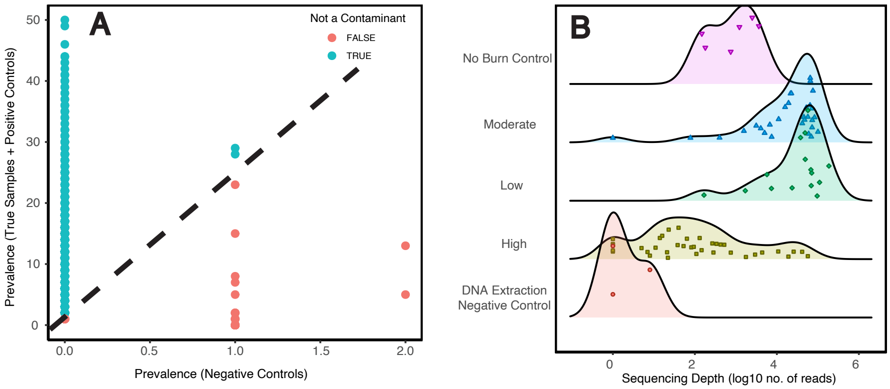


**Supplementary Fig. 5** Example assessment of procedures for removing contaminating DNA sequence reads —with Decontam [17] —from samples collected at the Decker Fire in November, 2019. These samples were the first samples collected after fire and are both low-biomass and likely contain high concentrations of PCR inhibitors. These types of samples are difficult to extract DNA from and assessing contamination is challenging. (A) Plotting the presence / absence (prevalence) of amplicon sequence variants (ASVs), defined as the number of samples that an individual ASV is present in, as a function of true samples vs. DNA extraction controls. The populations of classified contaminating vs. non-contaminating ASVs are separable (illustrated dashed line). (B) The sequencing depths of samples, by soil burn severity (SBS; as High, Moderate, Low, and No Burn), after identified contaminating reads have been removed. Peak heights represent sample density in the distribution (a histogram of samples as points). Populations of ASV sequencing depth are distinguishable by SBS, and all SBS classes have distributions of reads that are distinct from DNA extraction negative controls. Reads remaining in extraction controls after the removal of contaminants are ASVs that are also greatly present in the natural environment. All ASVs that passed the contamination procedure were retained for downstream analyses.

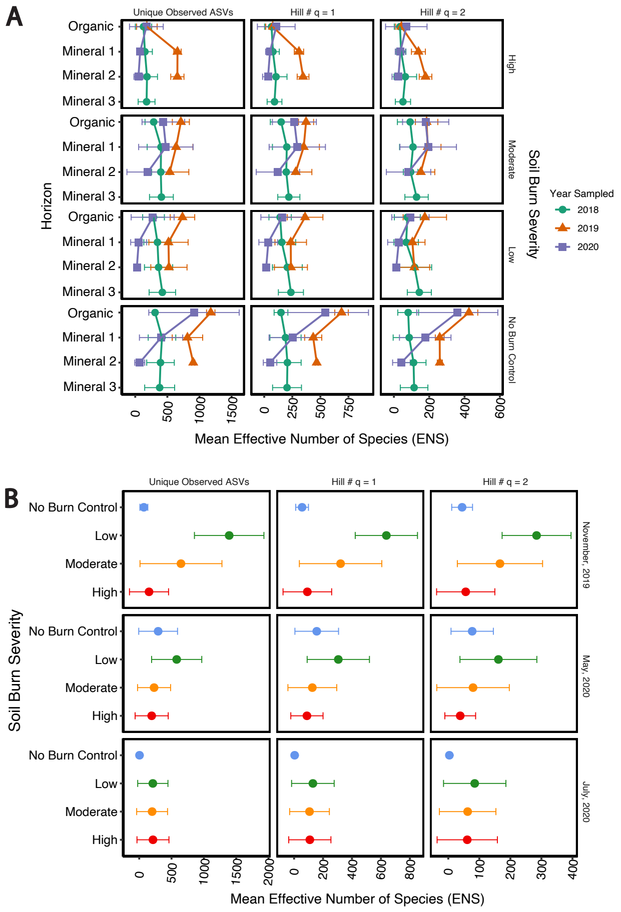


**Supplementary Fig. 6** Bacteria and Archaea microbial diversity after wildfire as measured by Unique Observed Amplicon Sequence Variants (ASVs) and Hill numbers (q = 1 and q = 2) +/- one standard deviation. In order, Observed ASVs, q = 1 Hill number, and q = 2 Hill number increase the weight of common microbiota in determining the effective number of species (ENS, x-axis) [19, 20]. With increasing weight on common species (increasing Hill #), the calculated ENS decreases. Soil microbial diversity fluctuates across all metadata factors. (A) The 416 Fire; samples were collected across soil burn severity (SBS), soil horizon, and time. (B) The Decker Fire; only organic horizon (Org-Horizon) samples were collected across SBS and time.

**Supplementary Table 1** Mean biogeochemical measurements +/- one standard deviation across replicate plots within our experimental design at the 416 Fire (soil burn severity, year, and soil horizon); N = 88. This is the first of three tables for the 416 Fire (see Supplementary Tables 2 and 3). A standard deviation of "NA" indicates that no replicate plots were measured.

| Soil Burn Severity | Soil Horizon | pH | ANC (ueq / L) | EC (uS) | DOC (mg / L) | DTN (mg / L) | Na (mg / L) |
|---|---|---|---|---|---|---|---|
| 2018 | | | | | | | |
| High | Organic | 8.55 +/- 0.18 | 7921 +/- 4016 | 1386 +/- 689 | 453.6 +/- 133.7 | 31.6 +/- 3.6 | 1.64 +/- 0.63 |
| High | Mineral 1 | 8.56 +/- 0.35 | 7320 +/- 3052 | 1056 +/- 430 | 276.8 +/- 202.0 | 26.2 +/- 12.2 | 1.38 +/- 0.39 |
| High | Mineral 2 | 8.44 +/- 0.75 | 4759 +/- 1228 | 720 +/- 88 | 147.3 +/- 113.8 | 15.4 +/- 9.1 | 1.18 +/- 0.04 |
| High | Mineral 3 | 8.21 +/- 0.32 | 4883 +/- 2632 | 713 +/- 302 | 149.2 +/- 112.1 | 13.9 +/- 12.8 | 1.16 +/- 0.58 |
| Moderate | Organic | 7.12 +/- 0.24 | 3974 +/- 4908 | 456 +/- 500 | 489.2 +/- 535.2 | 47.2 +/- 50.9 | 0.24 +/- 0.03 |
| Moderate | Mineral 1 | 7.08 +/- 0.26 | 1593 +/- 1466 | 185 +/- 142 | 48.2 +/- 34.1 | 3.4 +/- 1.5 | 0.27 +/- 0.01 |
| Moderate | Mineral 2 | 7.01 +/- 0.03 | 522 +/- 200 | 88 +/- 5 | 17.2 +/- 5.2 | 1.7 +/- 0.2 | 0.23 +/- 0.05 |
| Moderate | Mineral 3 | 6.99 +/- 0.2 | 704 +/- 102 | 94 +/- 23 | 16.3 +/- 1.3 | 1.6 +/- 0.2 | 0.15 +/- 0.01 |
| Low | Organic | 7.08 +/- 0.02 | 6971 +/- 4185 | 689 +/- 469 | 123.0 +/- 140.5 | 21.5 +/- 26.2 | 0.73 +/- 0.73 |
| Low | Mineral 1 | 6.89 +/- 0.05 | 5161 +/- 1486 | 556 +/- 217 | 121.3 +/- 148.7 | 15.6 +/- 19.7 | 0.34 +/- 0.23 |
| Low | Mineral 2 | 6.93 +/- 0.08 | 4055 +/- 605 | 415 +/- 17 | 20.3 +/- 4.7 | 2.8 +/- 1.5 | 0.27 +/- 0.11 |
| Low | Mineral 3 | 6.95 +/- 0.02 | 4139 +/- 431 | 412 +/- 8 | 18.5 +/- 5.9 | 1.9 +/- 0.2 | 0.19 +/- 0.02 |
| No Burn Control | Organic | 6.58 +/- NA | 1137 +/- NA | 133 +/- NA | 22.2 +/- NA | 3.8 +/- NA | 0.2 +/- NA |
| No Burn Control | Mineral 1 | 6.58 +/- NA | 1026 +/- NA | 134 +/- NA | 22.3 +/- NA | 2.9 +/- NA | 0.15 +/- NA |
| No Burn Control | Mineral 2 | 6.67 +/- NA | 721 +/- NA | 92 +/- NA | 19.5 +/- NA | 2.9 +/- NA | 0.15 +/- NA |
| No Burn Control | Mineral 3 | 6.72 +/- NA | 1095 +/- NA | 125 +/- NA | 19.7 +/- NA | 3.5 +/- NA | 0.16 +/- NA |
| 2019 | | | | | | | |
| High | Organic | 8.25 +/- 0.21 | 1174 +/- 857 | 188 +/- 157 | 33.6 +/- 17.9 | 7.3 +/- 5.2 | 0.55 +/- 0.38 |
| High | Mineral 1 | 8.16 +/- 0.37 | 415 +/- 287 | 63 +/- 37 | 12.2 +/- 2.3 | 1.5 +/- 0.1 | 0.49 +/- 0.35 |
| High | Mineral 2 | 7.87 +/- 0.74 | 300 +/- 140 | 47 +/- 19 | 10 +/- 1.0 | 1.4 +/- 0.3 | 0.3 +/- 0.06 |
| Moderate | Organic | 7.23 +/- 0.85 | 551 +/- 301 | 64.4 +/- 34.4 | 14.0 +/- 1.3 | 1.7 +/- 0.8 | 0.46 +/- 0.56 |
| Moderate | Mineral 1 | 7.15 +/- 0.6 | 418 +/- 254 | 52 +/- 24 | 11.3 +/- 2.4 | 1.5 +/- 0.5 | 0.5 +/- 0.52 |
| Moderate | Mineral 2 | 7.11 +/- 0.54 | 356 +/- 203 | 44 +/- 23 | 9.8 +/- 0.9 | 1.0 +/- 0.0 | 0.53 +/- 0.53 |
| Low | Organic | 7.13 +/- 0.76 | 484 +/- 488 | 54 +/- 51 | 16.7 +/- 6.4 | 1.0 +/- 0.1 | 1.0 +/- 0.4 |
| Low | Mineral 1 | 7.19 +/- 0.82 | 369 +/- 246 | 43 +/- 28 | 13.8 +/- 2.0 | 1.0 +/- 0.2 | 1.06 +/- 0.76 |
| Low | Mineral 2 | 7.32 +/- 0.87 | 220 +/- 56 | 28 +/- 7 | 10.5 +/- 3.0 | 0.9 +/- 0.2 | 0.96 +/- 0.66 |
| No Burn Control | Organic | 6.57 +/- NA | 667 +/- NA | 75 +/- NA | 21.2 +/- NA | 2.0 +/- NA | 0.13 +/- NA |
| No Burn Control | Mineral 1 | 6.31 +/- NA | 1064 +/- NA | 105 +/- NA | 22.1 +/- NA | 1.1 +/- NA | 0.14 +/- NA |
| No Burn Control | Mineral 2 | 6.42 +/- NA | 1069 +/- NA | 116 +/- NA | 21.2 +/- NA | 1.0 +/- NA | 0.14 +/- NA |
| 2020 | | | | | | | |
| High | Organic | 7.67 +/- 0.44 | 829 +/- 448 | 106 +/- 57 | 21.7 +/- 15.7 | 3.4 +/- 2.7 | 0.49 +/- 0.33 |
| High | Mineral 1 | 8.16 +/- 0.49 | 1068 +/- 383 | 135 +/- 70 | 24.7 +/- 25.3 | 4.1 +/- 3.6 | 0.56 +/- 0.39 |
| High | Mineral 2 | 7.98 +/- 0.54 | 649 +/- 62 | 79 +/- 2 | 7.6 +/- 4.1 | 1.2 +/- 0.0 | 0.59 +/- 0.22 |
| Moderate | Organic | 7.3 +/- 0.65 | 687 +/- 343 | 80 +/- 35 | 20.9 +/- 5.2 | 2.0 +/- 0.9 | 0.45 +/- 0.57 |
| Moderate | Mineral 1 | 7.14 +/- 0.57 | 535 +/- 280 | 59 +/- 30 | 10.4 +/- 4.3 | 1.0 +/- 0.2 | 0.5 +/- 0.53 |
| Moderate | Mineral 2 | 7.16 +/- 0.56 | 419 +/- 111 | 50 +/- 11 | 9.7 +/- 1.9 | 1.1 +/- 0.4 | 0.58 +/- 0.53 |
| Low | Organic | 7.26 +/- 0.71 | 1019 +/- 775 | 116 +/- 84 | 18.6 +/- 4.0 | 2.0 +/- 0.9 | 0.98 +/- 0.44 |
| Low | Mineral 1 | 7.2 +/- 0.66 | 624 +/- 369 | 67 +/- 36 | 25.0 +/- 19.1 | 1.6 +/- 0.6 | 0.94 +/- 0.54 |
| Low | Mineral 2 | 7.26 +/- 0.8 | 450 +/- 304 | 52 +/- 31 | 13.2 +/- 4.3 | 1.0 +/- 0.3 | 0.9 +/- 0.61 |
| No Burn Control | Organic | 6.79 +/- NA | 479 +/- NA | 65 +/- NA | 14.9 +/- NA | 1.5 +/- NA | 0.16 +/- NA |
| No Burn Control | Mineral 1 | 6.93 +/- NA | 795 +/- NA | 77 +/- NA | 9.8 +/- NA | 0.9 +/- NA | 0.18 +/- NA |
| No Burn Control | Mineral 2 | 6.94 +/- NA | 650 +/- NA | 65 +/- NA | 15.0 +/- NA | 1.0 +/- NA | 0.21 +/- NA |

Abbreviations: acid neutralizing capacity (ANC), electrical conductivity (EC), dissolved organic carbon (DOC), dissolved total nitrogen (DTN).

**Supplementary Table 2** Mean biogeochemical measurements +/- one standard deviation across replicate plots within our experimental design at the 416 Fire (soil burn severity, year, and soil horizon); N = 88. This is the second of three tables for the 416 Fire (see Supplementary Tables 1 and 3). A standard deviation of "NA" indicates that no replicate plots were measured.

| Soil Burn Severity | Soil Horizon | NH4 (mg / L) | K (mg / L) | Mg (mg / L) | Ca (mg / L) | Cl (mg / L) | NO3 (mg / L) |
|---|---|---|---|---|---|---|---|
| 2018 | | | | | | | |
| High | Organic | 2.01 +/- 1.08 | 308.55 +/- 117.31 | 16.92 +/- 23.44 | 30.2 +/- 13.96 | 19.02 +/- 16.54 | 56.57 +/- 45.9 |
| High | Mineral 1 | 2.04 +/- 1.36 | 294.4 +/- 92.35 | 12.9 +/- 17.46 | 23.88 +/- 7.31 | 18.1 +/- 14.35 | 48.79 +/- 57.14 |
| High | Mineral 2 | 4.45 +/- 4.77 | 189.05 +/- 39.81 | 5.92 +/- 7.68 | 18.77 +/- 3.44 | 6.13 +/- 0.9 | 9.66 +/- 7.25 |
| High | Mineral 3 | 1.87 +/- 1.54 | 185.1 +/- 134.21 | 7.96 +/- 10.72 | 17.24 +/- 0.52 | 5.11 +/- 0.09 | 3.79 +/- 0.6 |
| Moderate | Organic | 9.23 +/- 10.26 | 12.38 +/- 3.31 | 9.42 +/- 12.35 | 64.22 +/- 78.94 | 0.64 +/- 0.16 | 1.34 +/- 0.26 |
| Moderate | Mineral 1 | 0.35 +/- 0.24 | 13.31 +/- 2.08 | 1.08 +/- 0.14 | 28.28 +/- 30.96 | 0.58 +/- 0.36 | 1 +/- 0.38 |
| Moderate | Mineral 2 | 0.21 +/- 0.14 | 15.18 +/- 2.09 | 0.99 +/- 0.39 | 4.27 +/- 0.79 | 0.59 +/- 0.17 | 1.01 +/- 0.11 |
| Moderate | Mineral 3 | 0.2 +/- 0.13 | 16.14 +/- 5.23 | 0.87 +/- 0.51 | 5.84 +/- 0.51 | 0.51 +/- 0.15 | 1.43 +/- 0.88 |
| Low | Organic | 13.36 +/- 18.02 | 26.12 +/- 1.87 | 11.66 +/- 1.07 | 68.45 +/- 41.44 | 1.35 +/- 0.08 | 4.25 +/- 3.31 |
| Low | Mineral 1 | 7.95 +/- 11.02 | 24.34 +/- 1.44 | 13.12 +/- 3.16 | 71.04 +/- 29.67 | 1.36 +/- 0.19 | 3.48 +/- 2.52 |
| Low | Mineral 2 | 0.44 +/- 0.61 | 27.71 +/- 3.32 | 12.5 +/- 2.63 | 51.25 +/- 6.92 | 1.36 +/- 0.19 | 3.14 +/- 2.25 |
| Low | Mineral 3 | 0.17 +/- 0.23 | 25.11 +/- 1.23 | 11.88 +/- 1.86 | 53.06 +/- 1.76 | 1.27 +/- 0.01 | 1.53 +/- 0.15 |
| No Burn Control | Organic | 0.13 +/- NA | 29.84 +/- NA | 0.98 +/- NA | 7.09 +/- NA | 0.36 +/- NA | 3.62 +/- NA |
| No Burn Control | Mineral 1 | 0.18 +/- NA | 31.7 +/- NA | 0.74 +/- NA | 4.6 +/- NA | 0.33 +/- NA | 3.78 +/- NA |
| No Burn Control | Mineral 2 | 0.13 +/- NA | 20.4 +/- NA | 0.56 +/- NA | 3.1 +/- NA | 0.36 +/- NA | 3 +/- NA |
| No Burn Control | Mineral 3 | 0.12 +/- NA | 32.8 +/- NA | 0.51 +/- NA | 2.46 +/- NA | 0.35 +/- NA | 2.5 +/- NA |
| 2019 | | | | | | | |
| High | Organic | 1.12 +/- 1.09 | 31.28 +/- 28.03 | 2.03 +/- 1.68 | 10.86 +/- 8.56 | 0.53 +/- 0.09 | 14.69 +/- 19.17 |
| High | Mineral 1 | 0.05 +/- 0.07 | 9.37 +/- 10.65 | 0.68 +/- 0.55 | 4.65 +/- 0.15 | 0.63 +/- 0.24 | 2.13 +/- 0.18 |
| High | Mineral 2 | 0.02 +/- 0.02 | 6.04 +/- 6.14 | 0.46 +/- 0.2 | 4.45 +/- 0.61 | 0.76 +/- 0.11 | 1.37 +/- 0.04 |
| Moderate | Organic | 0.14 +/- 0.1 | 8.55 +/- 4.85 | 0.56 +/- 0.31 | 5.9 +/- 4.43 | 0.38 +/- 0.17 | 2.41 +/- 3.14 |
| Moderate | Mineral 1 | 0.11 +/- 0.06 | 7.5 +/- 3.99 | 0.46 +/- 0.34 | 4.24 +/- 3.78 | 0.47 +/- 0.21 | 0.93 +/- 0.44 |
| Moderate | Mineral 2 | 0.08 +/- 0.05 | 7.29 +/- 4.22 | 0.43 +/- 0.35 | 3.13 +/- 2.39 | 0.47 +/- 0.29 | 0.94 +/- 0.52 |
| Low | Organic | 0.15 +/- 0.06 | 3.56 +/- 3.33 | 0.75 +/- 0.53 | 6.79 +/- 7.72 | 0.53 +/- 0.03 | 0.28 +/- 0.19 |
| Low | Mineral 1 | 0.12 +/- 0.15 | 2.22 +/- 1.15 | 0.74 +/- 0.49 | 4.95 +/- 3.8 | 0.58 +/- 0.02 | 0.29 +/- 0.15 |
| Low | Mineral 2 | 0.1 +/- 0.09 | 1.81 +/- 0.59 | 0.7 +/- 0.51 | 2.47 +/- 0.8 | 0.5 +/- 0.13 | 0.27 +/- 0.11 |
| No Burn Control | Organic | 0.12 +/- NA | 20.24 +/- NA | 0.53 +/- NA | 0.94 +/- NA | 0.55 +/- NA | 0.71 +/- NA |
| No Burn Control | Mineral 1 | 0.19 +/- NA | 16.8 +/- NA | 0.31 +/- NA | 13.12 +/- NA | 0.41 +/- NA | 0.63 +/- NA |
| No Burn Control | Mineral 2 | 0.1 +/- NA | 20.7 +/- NA | 0.31 +/- NA | 13.12 +/- NA | 0.41 +/- NA | 0.67 +/- NA |
| 2020 | | | | | | | |
| High | Organic | 0.16 +/- 0.17 | 15.79 +/- 7.62 | 1.61 +/- 1.05 | 9.37 +/- 4.57 | 0.6 +/- 0.05 | 4.85 +/- 5.5 |
| High | Mineral 1 | 0.45 +/- 0.53 | 11.86 +/- 11.45 | 1.89 +/- 1.21 | 15.66 +/- 1.74 | 0.46 +/- 0.04 | 6.34 +/- 3.79 |
| High | Mineral 2 | 0.05 +/- 0.03 | 7.98 +/- 6.68 | 1.23 +/- 0.11 | 8.29 +/- 5.88 | 0.52 +/- 0.02 | 1.89 +/- 0.91 |
| Moderate | Organic | 0.18 +/- 0.1 | 12.01 +/- 7.29 | 1.23 +/- 0.56 | 6.66 +/- 4.3 | 0.42 +/- 0.15 | 1.56 +/- 1.05 |
| Moderate | Mineral 1 | 0.11 +/- 0.06 | 11.21 +/- 7.63 | 0.56 +/- 0.27 | 4.38 +/- 2.2 | 0.48 +/- 0.22 | 0.61 +/- 0.39 |
| Moderate | Mineral 2 | 0.08 +/- 0.06 | 8.5 +/- 2.37 | 0.47 +/- 0.28 | 3.54 +/- 1.59 | 0.42 +/- 0.13 | 0.56 +/- 0.53 |
| Low | Organic | 0.09 +/- 0.04 | 8.67 +/- 6.25 | 1.1 +/- 0.86 | 16.3 +/- 13.77 | 0.57 +/- 0.22 | 1.7 +/- 2.05 |
| Low | Mineral 1 | 0.06 +/- 0.09 | 4.72 +/- 3.28 | 0.71 +/- 0.32 | 9.54 +/- 5.96 | 0.6 +/- 0.23 | 0.16 +/- 0.14 |
| Low | Mineral 2 | 0.05 +/- 0.08 | 3.96 +/- 2.23 | 0.63 +/- 0.42 | 5.73 +/- 4.77 | 0.55 +/- 0.25 | 0.24 +/- 0.18 |
| No Burn Control | Organic | 0.14 +/- NA | 14.62 +/- NA | 0.86 +/- NA | 1.28 +/- NA | 0.27 +/- NA | 0.12 +/- NA |
| No Burn Control | Mineral 1 | 0.14 +/- NA | 12.6 +/- NA | 0.64 +/- NA | 8.53 +/- NA | 0.37 +/- NA | 0.11 +/- NA |
| No Burn Control | Mineral 2 | 0.14 +/- NA | 13.53 +/- NA | 0.33 +/- NA | 5.23 +/- NA | 0.25 +/- NA | 0.08 +/- NA |

**Supplementary Table 3** Mean biogeochemical measurements +/- one standard deviation across replicate plots within our experimental design at the 416 Fire (soil burn severity, year, and soil horizon); N = 88. This is the third of three tables for the 416 Fire (see Supplementary Tables 1 and 2). A standard deviation of “NA” indicates that no replicate plots were measured.

| Soil Burn Severity | Soil Horizon | PO4 (mg / L) | SO4 (mg / L) | NO3.N (mg / L) | NH4.N (mg / L) | DIN (mg / L) | DON (mg / L) |
|---|---|---|---|---|---|---|---|
| 2018 | | | | | | | |
| High | Organic | 3.15 +/- 1.32 | 58.53 +/- 11.97 | 12.78 +/- 10.37 | 1.56 +/- 0.84 | 14.34 +/- 9.53 | 17.3 +/- 6.0 |
| High | Mineral 1 | 6.28 +/- 4.61 | 58.79 +/- 8.32 | 11.02 +/- 12.91 | 1.58 +/- 1.05 | 12.61 +/- 11.85 | 13.5 +/- 0.4 |
| High | Mineral 2 | 3.75 +/- 0.61 | 52.96 +/- 22.5 | 2.18 +/- 1.64 | 3.46 +/- 3.7 | 5.64 +/- 2.06 | 9.7 +/- 11.1 |
| High | Mineral 3 | 2.36 +/- 0.37 | 51.54 +/- 19.48 | 0.86 +/- 0.14 | 1.45 +/- 1.19 | 2.31 +/- 1.33 | 11.6 +/- 14.1 |
| Moderate | Organic | 14.6 +/- 14.35 | 10.6 +/- 7.16 | 0.3 +/- 0.06 | 7.16 +/- 7.97 | 7.47 +/- 8.02 | 39.8 +/- 42.9 |
| Moderate | Mineral 1 | 2.52 +/- 0.11 | 5.4 +/- 0.82 | 0.23 +/- 0.09 | 0.27 +/- 0.18 | 0.5 +/- 0.1 | 2.9 +/- 1.4 |
| Moderate | Mineral 2 | 1.41 +/- 0.1 | 8.36 +/- 4.18 | 0.23 +/- 0.02 | 0.17 +/- 0.11 | 0.39 +/- 0.13 | 1.3 +/- 0.3 |
| Moderate | Mineral 3 | 1.95 +/- 1 | 4.25 +/- 0.12 | 0.32 +/- 0.2 | 0.16 +/- 0.1 | 0.48 +/- 0.3 | 1.1 +/- 0.1 |
| Low | Organic | 13.68 +/- 11.22 | 15.41 +/- 19.66 | 0.96 +/- 0.75 | 10.37 +/- 13.99 | 11.33 +/- 14.74 | 10.2 +/- 11.4 |
| Low | Mineral 1 | 14.24 +/- 12.38 | 13.86 +/- 17.9 | 0.79 +/- 0.57 | 6.17 +/- 8.56 | 6.96 +/- 9.12 | 8.7 +/- 10.6 |
| Low | Mineral 2 | 4.61 +/- 0.03 | 2.71 +/- 1.09 | 0.71 +/- 0.51 | 0.34 +/- 0.47 | 1.05 +/- 0.98 | 1.7 +/- 0.5 |
| Low | Mineral 3 | 4.56 +/- 0.4 | 2.56 +/- 1.57 | 0.35 +/- 0.03 | 0.13 +/- 0.18 | 0.48 +/- 0.15 | 1.5 +/- 0.4 |
| No Burn Control | Organic | 2.03 +/- NA | 1.17 +/- NA | 0.82 +/- NA | 0.1 +/- NA | 0.91 +/- NA | 2.9 +/- NA |
| No Burn Control | Mineral 1 | 1.53 +/- NA | 0.84 +/- NA | 0.85 +/- NA | 0.14 +/- NA | 0.99 +/- NA | 1.9 +/- NA |
| No Burn Control | Mineral 2 | 1.99 +/- NA | 0.76 +/- NA | 0.68 +/- NA | 0.1 +/- NA | 0.78 +/- NA | 2.1 +/- NA |
| No Burn Control | Mineral 3 | 1.56 +/- NA | 0.93 +/- NA | 0.57 +/- NA | 0.09 +/- NA | 0.66 +/- NA | 2.9 +/- NA |
| 2019 | | | | | | | |
| High | Organic | 2.93 +/- 0.71 | 4.47 +/- 3.07 | 3.32 +/- 4.33 | 0.87 +/- 0.85 | 4.19 +/- 3.48 | 3.1 +/- 1.7 |
| High | Mineral 1 | 2.55 +/- 0.44 | 4.15 +/- 2.95 | 0.48 +/- 0.04 | 0.04 +/- 0.05 | 0.52 +/- 0.09 | 0.9 +/- 0 |
| High | Mineral 2 | 2.26 +/- 0.1 | 4.26 +/- 2.97 | 0.31 +/- 0.01 | 0.01 +/- 0.01 | 0.32 +/- 0 | 1.0 +/- 0.3 |
| Moderate | Organic | 1.58 +/- 1.19 | 1.11 +/- 0.53 | 0.54 +/- 0.71 | 0.11 +/- 0.08 | 0.66 +/- 0.78 | 1.0 +/- 0.3 |
| Moderate | Mineral 1 | 0.75 +/- 0.46 | 1.06 +/- 0.42 | 0.21 +/- 0.1 | 0.08 +/- 0.04 | 0.29 +/- 0.07 | 1.2 +/- 0.5 |
| Moderate | Mineral 2 | 0.79 +/- 0.58 | 1.2 +/- 0.66 | 0.21 +/- 0.12 | 0.07 +/- 0.04 | 0.28 +/- 0.1 | 0.8 +/- 0.1 |
| Low | Organic | 0.65 +/- 0.45 | 1.51 +/- 0.88 | 0.06 +/- 0.04 | 0.11 +/- 0.04 | 0.18 +/- 0.09 | 0.8 +/- 0.1 |
| Low | Mineral 1 | 0.63 +/- 0.39 | 1.37 +/- 0.7 | 0.07 +/- 0.03 | 0.09 +/- 0.11 | 0.16 +/- 0.14 | 0.8 +/- 0.1 |
| Low | Mineral 2 | 0.6 +/- 0.41 | 1.6 +/- 1.16 | 0.06 +/- 0.03 | 0.08 +/- 0.07 | 0.14 +/- 0.09 | 0.7 +/- 0.3 |
| No Burn Control | Organic | 1.86 +/- NA | 0.95 +/- NA | 0.16 +/- NA | 0.09 +/- NA | 0.25 +/- NA | 1.7 +/- NA |
| No Burn Control | Mineral 1 | 1.48 +/- NA | 0.89 +/- NA | 0.14 +/- NA | 0.15 +/- NA | 0.29 +/- NA | 0.8 +/- NA |
| No Burn Control | Mineral 2 | 1.22 +/- NA | 0.95 +/- NA | 0.15 +/- NA | 0.08 +/- NA | 0.23 +/- NA | 0.8 +/- NA |
| 2020 | | | | | | | |
| High | Organic | 3.3 +/- 1.26 | 4.2 +/- 1.72 | 1.09 +/- 1.24 | 0.13 +/- 0.13 | 1.22 +/- 1.37 | 2.2 +/- 1.3 |
| High | Mineral 1 | 2.16 +/- 0.72 | 4.03 +/- 3.15 | 1.43 +/- 0.86 | 0.35 +/- 0.41 | 1.78 +/- 1.27 | 2.3 +/- 2.3 |
| High | Mineral 2 | 2.16 +/- 1.06 | 3.89 +/- 3.18 | 0.43 +/- 0.2 | 0.04 +/- 0.02 | 0.47 +/- 0.23 | 0.7 +/- 0.3 |
| Moderate | Organic | 2.4 +/- 1.96 | 1.06 +/- 0.51 | 0.35 +/- 0.24 | 0.14 +/- 0.08 | 0.49 +/- 0.23 | 1.5 +/- 0.7 |
| Moderate | Mineral 1 | 1.05 +/- 0.56 | 0.9 +/- 0.5 | 0.14 +/- 0.09 | 0.09 +/- 0.04 | 0.22 +/- 0.05 | 0.8 +/- 0.2 |
| Moderate | Mineral 2 | 0.82 +/- 0.27 | 0.92 +/- 0.29 | 0.13 +/- 0.12 | 0.07 +/- 0.05 | 0.19 +/- 0.08 | 0.9 +/- 0.4 |
| Low | Organic | 1.78 +/- 1.64 | 1.51 +/- 0.85 | 0.38 +/- 0.46 | 0.07 +/- 0.03 | 0.45 +/- 0.49 | 1.6 +/- 0.5 |
| Low | Mineral 1 | 0.49 +/- 0.15 | 1.74 +/- 0.82 | 0.04 +/- 0.03 | 0.05 +/- 0.07 | 0.09 +/- 0.1 | 1.5 +/- 0.6 |
| Low | Mineral 2 | 0.41 +/- 0.19 | 1.47 +/- 0.89 | 0.05 +/- 0.04 | 0.04 +/- 0.06 | 0.09 +/- 0.1 | 0.9 +/- 0.2 |
| No Burn Control | Organic | 2.06 +/- NA | 0.87 +/- NA | 0.03 +/- NA | 0.11 +/- NA | 0.14 +/- NA | 1.3 +/- NA |
| No Burn Control | Mineral 1 | 1.88 +/- NA | 0.82 +/- NA | 0.02 +/- NA | 0.11 +/- NA | 0.13 +/- NA | 0.8 +/- NA |
| No Burn Control | Mineral 2 | 1.84 +/- NA | 0.9 +/- NA | 0.02 +/- NA | 0.11 +/- NA | 0.13 +/- NA | 0.9 +/- NA |

Abbreviations: nitrate-nitrogen (NO3.N), ammonium-nitrogen (NH4.N), dissolved inorganic nitrogen (DIN), dissolved organic nitrogen (DON).

**Supplementary Table 4** Mean biogeochemical measurements +/- one standard deviation across replicate plots within our experimental design at the Decker Fire (soil burn severity, and year); N = 90.

| Soil Burn Severity | Soil Horizon | pH | ANC (ueq / L) | EC (uS) | DOC (mg / L) | DTN (mg / L) | Na (mg / L) |
|---|---|---|---|---|---|---|---|
| November 2019 | | | | | | | |
| High | Organic | 8.4 +/- 0.57 | 3713 +/- 1526 | 475 +/- 147 | 47.0 +/- 33.7 | 6.9 +/- 4.1 | 0.87 +/- 0.64 |
| Moderate | Organic | 7.8 +/- 0.42 | 2860 +/- 1780 | 349 +/- 192 | 30.5 +/- 17.3 | 7.3 +/- 3.3 | 0.41 +/- 0.33 |
| Low | Organic | 7.77 +/- 0.23 | 1567 +/- 840 | 207 +/- 63 | 19.4 +/- 13.7 | 9.2 +/- 4.5 | 0.26 +/- 0.18 |
| No Burn Control | Organic | 7.49 +/- 0.08 | 93 +/- 49 | 21 +/- 8 | 5.4 +/- 3.9 | 1.5 +/- 0.7 | 0.22 +/- 0.03 |
| May 2020 | | | | | | | |
| High | Organic | 8.17 +/- 0.35 | 3385 +/- 1210 | 432 +/- 138 | 96.7 +/- 102.1 | 17.4 +/- 26.2 | 0.67 +/- 0.72 |
| Moderate | Organic | 7.82 +/- 0.43 | 2929 +/- 2091 | 354 +/- 204 | 57.9 +/- 43.2 | 9.6 +/- 5.7 | 0.44 +/- 0.34 |
| Low | Organic | 8.04 +/- 0.23 | 1856 +/- 743 | 225 +/- 55 | 52.3 +/- 52.5 | 7.4 +/- 5.5 | 0.29 +/- 0.15 |
| No Burn Control | Organic | 7.48 +/- 0.02 | 568 +/- 49 | 68 +/- 14 | 24.1 +/- 27.3 | 1.9 +/- 1.1 | 0.23 +/- 0.02 |
| July 2020 | | | | | | | |
| High | Organic | 8.19 +/- 0.34 | 3396 +/- 1290 | 430 +/- 151 | 77.1 +/- 51.6 | 8.5 +/- 3.9 | 0.67 +/- 0.68 |
| Moderate | Organic | 7.82 +/- 0.39 | 2795 +/- 1920 | 341 +/- 202 | 39.2 +/- 24.0 | 7.3 +/- 4.4 | 0.38 +/- 0.35 |
| Low | Organic | 8.03 +/- 0.25 | 1937 +/- 754 | 229 +/- 60 | 24.6 +/- 13.7 | 5.0 +/- 2.3 | 0.23 +/- 0.15 |
| No Burn Control | Organic | 7.48 +/- 0.01 | 461 +/- 21 | 47 +/- 6 | 2.3 +/- 0.1 | 0.6 +/- 0.0 | 0.17 +/- 0.08 |

| Soil Burn Severity | Soil Horizon | NH4 (mg / L) | K (mg / L) | Mg (mg / L) | Ca (mg / L) | Cl (mg / L) | NO3 (mg / L) |
|---|---|---|---|---|---|---|---|
| November 2019 | | | | | | | |
| High | Organic | 2.13 +/- 2.39 | 67.28 +/- 24.71 | 11.99 +/- 9.31 | 31.59 +/- 27.77 | 2.27 +/- 2.5 | 2.29 +/- 2.74 |
| Moderate | Organic | 3.74 +/- 3.52 | 42.83 +/- 28.54 | 7.44 +/- 7.33 | 28.86 +/- 30.69 | 0.66 +/- 0.31 | 5.83 +/- 7.59 |
| Low | Organic | 0.89 +/- 0.71 | 32.85 +/- 11.53 | 2.22 +/- 1.14 | 18.65 +/- 17.75 | 0.64 +/- 0.24 | 13.11 +/- 13.49 |
| No Burn Control | Organic | 0.32 +/- 0.08 | 3.13 +/- 1.56 | 0.25 +/- 0.2 | 1.21 +/- 0.38 | 0.45 +/- 0 | 3.43 +/- 1.57 |
| May 2020 | | | | | | | |
| High | Organic | 3.26 +/- 2.74 | 62.45 +/- 40.2 | 10.47 +/- 8.52 | 27.03 +/- 17.47 | 2.06 +/- 2.12 | 3.03 +/- 2.83 |
| Moderate | Organic | 4.77 +/- 4.77 | 47 +/- 45.67 | 7.74 +/- 7.83 | 28.59 +/- 35.15 | 0.59 +/- 0.4 | 4.82 +/- 4.25 |
| Low | Organic | 2.05 +/- 2.77 | 38.93 +/- 16.82 | 2.65 +/- 1.44 | 18.95 +/- 16.09 | 0.85 +/- 0.53 | 5.2 +/- 3.61 |
| No Burn Control | Organic | 0.41 +/- 0.44 | 6.82 +/- 3.79 | 0.87 +/- 0.03 | 7.65 +/- 0.45 | 0.25 +/- 0.16 | 2.59 +/- 2.7 |
| July 2020 | | | | | | | |
| High | Organic | 2.57 +/- 2.34 | 62.99 +/- 40.08 | 10.73 +/- 8.38 | 26.77 +/- 17.47 | 2.06 +/- 2.17 | 2.83 +/- 2.84 |
| Moderate | Organic | 2.86 +/- 4.01 | 43.28 +/- 36.21 | 8.12 +/- 8.72 | 28.81 +/- 36.17 | 0.5 +/- 0.3 | 5.94 +/- 4.09 |
| Low | Organic | 0.75 +/- 0.81 | 39.14 +/- 17.28 | 2.44 +/- 1.17 | 19.95 +/- 17.16 | 0.81 +/- 0.5 | 5.3 +/- 3.01 |
| No Burn Control | Organic | 0.03 +/- 0.02 | 3.73 +/- 1.48 | 0.69 +/- 0.01 | 6.51 +/- 0 | 0.05 +/- 0.02 | 1.06 +/- 0.8 |

| Soil Burn Severity | Soil Horizon | PO4 (mg / L) | SO4 (mg / L) | NO3.N (mg / L) | NH4.N (mg / L) | DIN (mg / L) | DON (mg / L) |
|---|---|---|---|---|---|---|---|
| November 2019 | | | | | | | |
| High | Organic | 3.93 +/- 3.26 | 26.79 +/- 20.21 | 0.52 +/- 0.62 | 1.65 +/- 1.85 | 2.17 +/- 2.11 | 4.7 +/- 3.5 |
| Moderate | Organic | 5.73 +/- 3.05 | 8.24 +/- 4.75 | 1.32 +/- 1.71 | 2.91 +/- 2.74 | 4.22 +/- 2.49 | 3.1 +/- 3.3 |
| Low | Organic | 4.63 +/- 3.27 | 3.35 +/- 2.29 | 2.96 +/- 3.05 | 0.69 +/- 0.55 | 3.66 +/- 2.94 | 5.6 +/- 4.7 |
| No Burn Control | Organic | 1.03 +/- 0.42 | 0.58 +/- 0.11 | 0.78 +/- 0.35 | 0.25 +/- 0.06 | 1.03 +/- 0.29 | 0.4 +/- 0.4 |
| May 2020 | | | | | | | |
| High | Organic | 5.22 +/- 1.8 | 17.39 +/- 8.61 | 0.68 +/- 0.64 | 2.53 +/- 2.13 | 3.22 +/- 1.98 | 14.2 +/- 26.2 |
| Moderate | Organic | 6.01 +/- 2.26 | 8.04 +/- 7.79 | 1.09 +/- 0.96 | 3.7 +/- 3.71 | 4.79 +/- 3.69 | 4.8 +/- 4.4 |
| Low | Organic | 7.02 +/- 4.59 | 4.39 +/- 3.5 | 1.18 +/- 0.82 | 1.59 +/- 2.15 | 2.76 +/- 2.38 | 4.6 +/- 3.7 |
| No Burn Control | Organic | 3.4 +/- 3.08 | 0.71 +/- 0.77 | 0.59 +/- 0.61 | 0.31 +/- 0.35 | 0.9 +/- 0.27 | 1.0 +/- 0.8 |
| July 2020 | | | | | | | |
| High | Organic | 4.85 +/- 1.47 | 17.73 +/- 8.59 | 0.64 +/- 0.64 | 2 +/- 1.82 | 2.64 +/- 1.75 | 5.9 +/- 3.2 |
| Moderate | Organic | 5.9 +/- 2.14 | 8.35 +/- 8.61 | 1.34 +/- 0.92 | 2.22 +/- 3.12 | 3.57 +/- 2.9 | 3.7 +/- 4.1 |
| Low | Organic | 5.37 +/- 3.21 | 4.27 +/- 3.34 | 1.2 +/- 0.68 | 0.58 +/- 0.63 | 1.78 +/- 0.99 | 3.3 +/- 2.4 |
| No Burn Control | Organic | 2.02 +/- 2.01 | 0.19 +/- 0.11 | 0.24 +/- 0.18 | 0.02 +/- 0.02 | 0.26 +/- 0.16 | 0.3 +/- 0.2 |

Abbreviations: acid neutralizing capacity (ANC), electrical conductivity (EC), dissolved organic carbon (DOC), dissolved total nitrogen (DTN), nitrate-nitrogen (NO3.N), ammonium-nitrogen (NH4.N), dissolved inorganic nitrogen (DIN), dissolved organic nitrogen (DON).

**Supplementary Table 5** Summary statistics for the 13 measured biogeochemistry features that were fed into statistical learning models (N = 178 samples). All predictions in this manuscript were made by regressing on the log10-transformed target feature (except for pH since, by definition, it is already in log-space).

| | | Min. | 1st Qu. | Median | Mean | 3rd Qu. | Max. |
|---|---|---|---|---|---|---|---|
| **Normal Space** | DTN (mg / L) | 0.6 | 1.5 | 3.4 | 7.0 | 8.4 | 99.1 |
| | DOC (mg / L) | 2.2 | 12.5 | 22.2 | 52.4 | 46.2 | 867.6 |
| | ANC (ueq / L) | 58 | 603 | 1456 | 2186 | 3019 | 10760 |
| | PO4 (mg / L) | 0.19 | 1.34 | 2.88 | 3.92 | 5.40 | 24.74 |
| | SO4 (mg / L) | 0.11 | 1.21 | 4.23 | 9.81 | 11.76 | 84.47 |
| | K (mg / L) | 1.16 | 10.39 | 23.78 | 40.92 | 39.22 | 391.50 |
| | NH4 (mg / L) | 0.00 | 0.12 | 0.43 | 1.84 | 2.11 | 26.10 |
| | NO3 (mg / L) | 0.07 | 0.51 | 1.64 | 4.53 | 4.60 | 89.19 |
| | Ca (mg / L) | 0.94 | 4.92 | 12.39 | 19.95 | 21.28 | 124.75 |
| | Mg (mg / L) | 0.10 | 0.61 | 1.90 | 5.28 | 9.19 | 36.08 |
| | Na (mg / L) | 0.03 | 0.21 | 0.29 | 0.55 | 0.75 | 2.40 |
| | Cl (mg / L) | 0.04 | 0.42 | 0.62 | 1.42 | 0.99 | 30.72 |
| | pH | 6.31 | 6.95 | 7.90 | 7.66 | 8.09 | 9.69 |
| | | Min. | 1st Qu. | Median | Mean | 3rd Qu. | Max. |
| **log10 Transformed** | DTN (mg / L) | -0.23 | 0.16 | 0.54 | 0.85 | 0.93 | 2.00 |
| | DOC (mg / L) | 0.35 | 1.10 | 1.35 | 1.72 | 1.66 | 2.94 |
| | ANC (ueq / L) | 1.76 | 2.78 | 3.16 | 3.34 | 3.48 | 4.03 |
| | PO4 (mg / L) | -0.72 | 0.13 | 0.46 | 0.59 | 0.73 | 1.39 |
| | SO4 (mg / L) | -0.95 | 0.08 | 0.63 | 0.99 | 1.07 | 1.93 |
| | K (mg / L) | 0.07 | 1.02 | 1.38 | 1.61 | 1.59 | 2.59 |
| | NH4 (mg / L) | -3.00 | -0.93 | -0.37 | 0.26 | 0.32 | 1.42 |
| | NO3 (mg / L) | -1.16 | -0.29 | 0.22 | 0.66 | 0.66 | 1.95 |
| | Ca (mg / L) | -0.03 | 0.69 | 1.09 | 1.30 | 1.33 | 2.10 |
| | Mg (mg / L) | -0.99 | -0.21 | 0.28 | 0.72 | 0.96 | 1.56 |
| | Na (mg / L) | -1.47 | -0.68 | -0.54 | -0.26 | -0.13 | 0.38 |
| | Cl (mg / L) | -1.46 | -0.37 | -0.21 | 0.15 | -0.00 | 1.49 |

Abbreviations: dissolved total nitrogen (DTN), dissolved organic carbon (DOC), acid neutralizing capacity (ANC).

**Supplementary Table 6** Biogeochemistry-only based predictions of biogeochemical features after wildfire. All target features (except pH) were log10 transformed prior to modeling. Root mean squared error (RMSE) of residuals was used to evaluate models; RMSE measures how close predictions are to the truth. The randomized set of training / testing data that produced the median testing RMSE (across 10 different randomized sets) for dissolved organic carbon (DOC) and dissolved total nitrogen (DTN) was used for all of the predictions shown here. The number of learned coefficients selected from biogeochemistry features are reported; the intercept term was included in all models.

| | | Training RMSE | Testing RMSE | # Biogeochem. Features | # Microbio. Features | Fraction from Rare Biosphere |
|---|---|---|---|---|---|---|
| **Biogeochem. Only** | DTN | 0.25 | 0.32 | 5 | 0 | NA |
| | DOC | 0.29 | 0.29 | 4 | 0 | NA |
| | ANC | 0.20 | 0.19 | 10 | 0 | NA |
| | PO4 | 0.34 | 0.34 | 8 | 0 | NA |
| | SO4 | 0.33 | 0.41 | 8 | 0 | NA |
| | K | 0.31 | 0.32 | 11 | 0 | NA |
| | NH4 | 0.66 | 0.74 | 6 | 0 | NA |
| | NO3 | 0.55 | 0.39 | 5 | 0 | NA |
| | Ca | 0.28 | 0.24 | 3 | 0 | NA |
| | Mg | 0.38 | 0.29 | 8 | 0 | NA |
| | Na | 0.30 | 0.36 | 3 | 0 | NA |
| | Cl | 0.22 | 0.29 | 10 | 0 | NA |
| | pH | 0.58 | 0.57 | 2 | 0 | NA |

NA indicates Not Applicable. Abbreviations: dissolved total nitrogen (DTN), dissolved organic carbon (DOC), acid neutralizing capacity (ANC).

**Supplementary Table 7** Hybrid biogeochemistry-microbiome based predictions of biogeochemical features after wildfire. Models at the Phylum, Class, and Order taxonomic ranks are shown. All target features (except pH) were log10 transformed prior to modeling. Root mean squared error (RMSE) of residuals was used to evaluate models; RMSE measures how close predictions are to the truth. The randomized set of training / testing data that produced the median testing RMSE (across 10 different randomized sets) for dissolved organic carbon (DOC) and dissolved total nitrogen (DTN) was used for all of the predictions shown here. The number of learned coefficients selected from each of biogeochemistry and the microbiome are reported; the intercept term was included in all models. A taxon of microorganism is defined as being from the rare biosphere if it has a mean relative abundance of $< 1\%$.

| | | Training RMSE | Testing RMSE | # Biogeochem. Features | # Microbio. Features | Fraction from Rare Biosphere |
|---|---|---|---|---|---|---|
| Phylum | DTN | 0.25 | 0.27 | 6 | 3 | 0.67 |
| | DOC | 0.22 | 0.35 | 7 | 23 | 0.78 |
| | ANC | 0.20 | 0.35 | 8 | 11 | 0.64 |
| | PO4 | 0.32 | 0.40 | 6 | 10 | 0.60 |
| | SO4 | 0.30 | 0.35 | 7 | 28 | 0.71 |
| | K | 0.29 | 0.50 | 7 | 12 | 0.75 |
| | NH4 | 0.54 | 0.76 | 7 | 13 | 0.62 |
| | NO3 | 0.46 | 0.49 | 5 | 25 | 0.68 |
| | Ca | 0.24 | 0.58 | 5 | 12 | 0.75 |
| | Mg | 0.26 | 0.46 | 11 | 44 | 0.80 |
| | Na | 0.28 | 0.37 | 3 | 8 | 0.38 |
| | Cl | 0.20 | 0.28 | 9 | 30 | 0.80 |
| | pH | 0.46 | 0.68 | 5 | 22 | 0.73 |
| Class | DTN | 0.25 | 0.27 | 6 | 3 | 0.33 |
| | DOC | 0.22 | 0.24 | 6 | 19 | 0.84 |
| | ANC | 0.19 | 2.02 | 7 | 18 | 0.83 |
| | PO4 | 0.32 | 0.30 | 6 | 10 | 0.70 |
| | SO4 | 0.31 | 0.79 | 6 | 31 | 0.74 |
| | K | 0.25 | 1.97 | 7 | 35 | 0.83 |
| | NH4 | 0.49 | 0.77 | 7 | 30 | 0.83 |
| | NO3 | 0.49 | 0.46 | 4 | 15 | 0.73 |
| | Ca | 0.22 | 0.82 | 5 | 35 | 0.86 |
| | Mg | 0.32 | 0.42 | 6 | 31 | 0.87 |
| | Na | 0.28 | 0.41 | 3 | 7 | 0.57 |
| | Cl | 0.21 | 0.26 | 7 | 15 | 0.73 |
| | pH | 0.45 | 0.52 | 5 | 19 | 0.84 |
| Order | DTN | 0.25 | 0.26 | 6 | 5 | 0.60 |
| | DOC | 0.23 | 0.24 | 4 | 12 | 0.58 |
| | ANC | 0.16 | 0.23 | 8 | 47 | 0.87 |
| | PO4 | 0.32 | 0.31 | 5 | 8 | 0.75 |
| | SO4 | 0.32 | 0.34 | 6 | 10 | 0.50 |
| | K | 0.29 | 0.35 | 5 | 24 | 0.75 |
| | NH4 | 0.48 | 0.59 | 6 | 39 | 0.82 |
| | NO3 | 0.43 | 0.46 | 5 | 45 | 0.84 |
| | Ca | 0.20 | 0.27 | 5 | 68 | 0.99 |
| | Mg | 0.28 | 0.41 | 7 | 53 | 0.87 |
| | Na | 0.28 | 0.39 | 2 | 13 | 0.77 |
| | Cl | 0.20 | 0.24 | 8 | 28 | 0.86 |
| | pH | 0.49 | 0.50 | 4 | 20 | 0.80 |

Abbreviations: dissolved total nitrogen (DTN), dissolved organic carbon (DOC), acid neutralizing capacity (ANC).

**Supplementary Table 8** Hybrid biogeochemistry-microbiome based predictions of biogeochemical features after wildfire. Models at the Family, Genus, and ASV taxonomic ranks are shown. All target features (except pH) were log10 transformed prior to modeling. Root mean squared error (RMSE) of residuals was used to evaluate models; RMSE measures how close predictions are to the truth. The randomized set of training / testing data that produced the median testing RMSE (across 10 different randomized sets) for dissolved organic carbon (DOC) and dissolved total nitrogen (DTN) was used for all of the predictions shown here. The number of learned coefficients selected from each of biogeochemistry and the microbiome are reported; the intercept term was included in all models. A taxon of microorganism is defined as being from the rare biosphere if it has a mean relative abundance of $< 1\%$.

| | | Training RMSE | Testing RMSE | # Biogeochem. Features | # Microbio. Features | Fraction from Rare Biosphere |
|---|---|---|---|---|---|---|
| Family | DTN | 0.23 | 0.25 | 6 | 6 | 0.83 |
| | DOC | 0.21 | 0.23 | 4 | 38 | 0.87 |
| | ANC | 0.16 | 0.21 | 8 | 40 | 0.85 |
| | PO4 | 0.31 | 0.29 | 5 | 17 | 0.76 |
| | SO4 | 0.30 | 0.32 | 6 | 21 | 0.71 |
| | K | 0.23 | 0.40 | 7 | 55 | 0.84 |
| | NH4 | 0.40 | 0.65 | 7 | 58 | 0.88 |
| | NO3 | 0.49 | 0.46 | 4 | 19 | 0.79 |
| | Ca | 0.22 | 0.26 | 3 | 46 | 0.96 |
| | Mg | 0.35 | 0.43 | 4 | 27 | 0.89 |
| | Na | 0.26 | 0.39 | 2 | 20 | 0.90 |
| | Cl | 0.22 | 0.25 | 6 | 12 | 0.92 |
| | pH | 0.49 | 0.49 | 4 | 19 | 0.74 |
| Genus | DTN | 0.21 | 0.23 | 6 | 15 | 0.80 |
| | DOC | 0.21 | 0.24 | 4 | 24 | 0.75 |
| | ANC | 0.11 | 0.18 | 8 | 109 | 0.94 |
| | PO4 | 0.30 | 0.28 | 5 | 16 | 0.81 |
| | SO4 | 0.29 | 0.33 | 6 | 39 | 0.79 |
| | K | 0.24 | 0.32 | 6 | 53 | 0.85 |
| | NH4 | 0.45 | 0.60 | 6 | 50 | 0.86 |
| | NO3 | 0.36 | 0.46 | 6 | 91 | 0.97 |
| | Ca | 0.24 | 0.26 | 4 | 32 | 0.97 |
| | Mg | 0.38 | 0.44 | 4 | 18 | 0.83 |
| | Na | 0.25 | 0.36 | 2 | 35 | 0.97 |
| | Cl | 0.22 | 0.25 | 6 | 17 | 0.88 |
| | pH | 0.40 | 0.47 | 4 | 52 | 0.96 |
| ASV | DTN | 0.26 | 0.28 | 6 | 2 | 0.50 |
| | DOC | 0.24 | 0.26 | 4 | 20 | 0.95 |
| | ANC | 0.13 | 0.19 | 7 | 80 | 1.00 |
| | PO4 | 0.29 | 0.30 | 5 | 32 | 0.97 |
| | SO4 | 0.21 | 0.32 | 6 | 104 | 0.97 |
| | K | 0.30 | 0.36 | 4 | 22 | 1.00 |
| | NH4 | 0.36 | 0.56 | 6 | 109 | 0.97 |
| | NO3 | 0.46 | 0.43 | 4 | 28 | 0.96 |
| | Ca | 0.23 | 0.26 | 4 | 40 | 1.00 |
| | Mg | 0.14 | 0.43 | 7 | 226 | 1.00 |
| | Na | 0.25 | 0.31 | 2 | 38 | 1.00 |
| | Cl | 0.23 | 0.28 | 7 | 11 | 1.00 |
| | pH | 0.20 | 0.39 | 3 | 205 | 1.00 |

Abbreviations: dissolved total nitrogen (DTN), dissolved organic carbon (DOC), acid neutralizing capacity (ANC).

**Supplementary Table 9** Microbiome-only based predictions of biogeochemical features after wildfire. Models at the Phylum, Class, and Order taxonomic ranks are shown. All target features (except pH) were log10 transformed prior to modeling. Root mean squared error (RMSE) of residuals was used to evaluate models; RMSE measures how close predictions are to the truth. The randomized set of training / testing data that produced the median testing RMSE (across 10 different randomized sets) for dissolved organic carbon (DOC) and dissolved total nitrogen (DTN) was used for all of the predictions shown here. The number of learned coefficients selected from the microbiome are reported; the intercept term was included in all models. A taxon of microorganism is defined as being from the rare biosphere if it has a mean relative abundance of < 1%.

| | | Training RMSE | Testing RMSE | # Biogeochem. Features | # Microbio. Features | Fraction from Rare Biosphere |
|---|---|---|---|---|---|---|
| Phylum | DTN | 0.41 | 0.47 | 0 | 4 | 0.50 |
| | DOC | 0.39 | 0.40 | 0 | 16 | 0.56 |
| | ANC | 0.40 | 0.66 | 0 | 13 | 0.62 |
| | PO4 | 0.38 | 0.55 | 0 | 12 | 0.58 |
| | SO4 | 0.49 | 0.54 | 0 | 20 | 0.70 |
| | K | 0.43 | 0.73 | 0 | 15 | 0.67 |
| | NH4 | 0.73 | 0.94 | 0 | 6 | 0.33 |
| | NO3 | 0.57 | 0.58 | 0 | 14 | 0.57 |
| | Ca | 0.42 | 0.93 | 0 | 14 | 0.79 |
| | Mg | 0.53 | 0.62 | 0 | 19 | 0.68 |
| | Na | 0.32 | 0.39 | 0 | 12 | 0.75 |
| | Cl | 0.37 | 0.43 | 0 | 8 | 0.50 |
| | pH | 0.54 | 0.56 | 0 | 31 | 0.77 |
| Class | DTN | 0.40 | 0.45 | 0 | 12 | 0.50 |
| | DOC | 0.33 | 0.41 | 0 | 35 | 0.80 |
| | ANC | 0.38 | 0.45 | 0 | 17 | 0.71 |
| | PO4 | 0.40 | 0.42 | 0 | 6 | 0.50 |
| | SO4 | 0.53 | 0.57 | 0 | 14 | 0.50 |
| | K | 0.43 | 0.52 | 0 | 14 | 0.64 |
| | NH4 | 0.68 | 0.90 | 0 | 13 | 0.69 |
| | NO3 | 0.64 | 0.59 | 0 | 0 | NA |
| | Ca | 0.47 | 0.47 | 0 | 1 | 0.00 |
| | Mg | 0.49 | 0.58 | 0 | 35 | 0.83 |
| | Na | 0.32 | 0.44 | 0 | 14 | 0.86 |
| | Cl | 0.35 | 3.85 | 0 | 20 | 0.80 |
| | pH | 0.57 | 0.57 | 0 | 18 | 0.83 |
| Order | DTN | 0.37 | 0.43 | 0 | 15 | 0.47 |
| | DOC | 0.38 | 0.43 | 0 | 12 | 0.42 |
| | ANC | 0.33 | 0.37 | 0 | 47 | 0.89 |
| | PO4 | 0.36 | 0.36 | 0 | 15 | 0.73 |
| | SO4 | 0.45 | 0.53 | 0 | 35 | 0.77 |
| | K | 0.42 | 0.49 | 0 | 18 | 0.67 |
| | NH4 | 0.50 | 0.73 | 0 | 69 | 0.86 |
| | NO3 | 0.48 | 0.58 | 0 | 59 | 0.86 |
| | Ca | 0.38 | 0.43 | 0 | 39 | 0.92 |
| | Mg | 0.54 | 0.57 | 0 | 17 | 0.65 |
| | Na | 0.33 | 0.39 | 0 | 4 | 0.75 |
| | Cl | 0.40 | 0.46 | 0 | 0 | NA |
| | pH | 0.56 | 0.57 | 0 | 22 | 0.77 |

NA indicates Not Applicable. Abbreviations: dissolved total nitrogen (DTN), dissolved organic carbon (DOC), acid neutralizing capacity (ANC).

**Supplementary Table 10** Microbiome-only based predictions of biogeochemical features after wildfire. Models at the Family, Genus, and ASV taxonomic ranks are shown. All target features (except pH) were log10 transformed prior to modeling. Root mean squared error (RMSE) of residuals was used to evaluate models; RMSE measures how close predictions are to the truth. The randomized set of training / testing data that produced the median testing RMSE (across 10 different randomized sets) for dissolved organic carbon (DOC) and dissolved total nitrogen (DTN) was used for all of the predictions shown here. The number of learned coefficients selected from the microbiome are reported; the intercept term was included in all models. A taxon of microorganism is defined as being from the rare biosphere if it has a mean relative abundance of $< 1\%$.

| | | Training RMSE | Testing RMSE | # Biogeochem. Features | # Microbio. Features | Fraction from Rare Biosphere |
|---|---|---|---|---|---|---|
| **Family** | DTN | 0.36 | 0.43 | 0 | 16 | 0.50 |
| | DOC | 0.36 | 0.41 | 0 | 23 | 0.70 |
| | ANC | 0.32 | 0.48 | 0 | 48 | 0.90 |
| | PO4 | 0.33 | 0.35 | 0 | 20 | 0.75 |
| | SO4 | 0.36 | 0.55 | 0 | 72 | 0.86 |
| | K | 0.38 | 0.47 | 0 | 32 | 0.78 |
| | NH4 | 0.50 | 0.69 | 0 | 66 | 0.88 |
| | NO3 | 0.60 | 0.56 | 0 | 19 | 0.68 |
| | Ca | 0.37 | 0.41 | 0 | 47 | 0.94 |
| | Mg | 0.48 | 0.61 | 0 | 41 | 0.85 |
| | Na | 0.31 | 0.40 | 0 | 14 | 0.93 |
| | Cl | 0.39 | 0.45 | 0 | 3 | 0.67 |
| | pH | 0.57 | 0.56 | 0 | 27 | 0.78 |
| **Genus** | DTN | 0.31 | 1.10 | 0 | 48 | 0.81 |
| | DOC | 0.32 | 0.45 | 0 | 35 | 0.74 |
| | ANC | 0.26 | 0.34 | 0 | 95 | 0.92 |
| | PO4 | 0.33 | 0.34 | 0 | 32 | 0.78 |
| | SO4 | 0.31 | 0.50 | 0 | 106 | 0.93 |
| | K | 0.34 | 0.43 | 0 | 52 | 0.87 |
| | NH4 | 0.54 | 0.72 | 0 | 57 | 0.84 |
| | NO3 | 0.56 | 0.54 | 0 | 22 | 0.82 |
| | Ca | 0.36 | 0.41 | 0 | 52 | 0.96 |
| | Mg | 0.50 | 0.57 | 0 | 43 | 0.91 |
| | Na | 0.31 | 0.41 | 0 | 28 | 0.96 |
| | Cl | 0.38 | 0.44 | 0 | 6 | 0.83 |
| | pH | 0.52 | 0.53 | 0 | 46 | 0.91 |
| **ASV** | DTN | 0.23 | 0.63 | 0 | 129 | 0.97 |
| | DOC | 0.29 | 0.41 | 0 | 72 | 0.96 |
| | ANC | 0.29 | 0.36 | 0 | 84 | 0.99 |
| | PO4 | 0.30 | 0.35 | 0 | 51 | 0.96 |
| | SO4 | 0.30 | 0.55 | 0 | 131 | 0.98 |
| | K | 0.15 | 0.40 | 0 | 214 | 0.99 |
| | NH4 | 0.43 | 0.73 | 0 | 121 | 0.98 |
| | NO3 | 0.33 | 0.52 | 0 | 150 | 0.99 |
| | Ca | 0.39 | 0.43 | 0 | 46 | 1.00 |
| | Mg | 0.23 | 0.46 | 0 | 218 | 0.99 |
| | Na | 0.31 | 0.37 | 0 | 25 | 1.00 |
| | Cl | 0.30 | 0.41 | 0 | 50 | 0.98 |
| | pH | 0.17 | 0.52 | 0 | 255 | 1.00 |

Abbreviations: dissolved total nitrogen (DTN), dissolved organic carbon (DOC), acid neutralizing capacity (ANC).
